# Whole-genome duplication in the Multicellularity Long Term Evolution Experiment

**DOI:** 10.1101/2024.04.18.588554

**Authors:** Kai Tong, Sayantan Datta, Vivian Cheng, Daniella J. Haas, Saranya Gourisetti, Harley L. Yopp, Thomas C. Day, Dung T. Lac, Peter L. Conlin, G. Ozan Bozdag, William C. Ratcliff

## Abstract

Whole-genome duplication (WGD) is widespread across eukaryotes and can promote adaptive evolution^1–4^. However, given the instability of newly-formed polyploid genomes^5–7^, understanding how WGDs arise in a population, persist, and underpin adaptations remains a challenge. Using our ongoing Multicellularity Long Term Evolution Experiment (MuLTEE)^8^, we show that diploid snowflake yeast (*Saccharomyces cerevisiae*) under selection for larger multicellular size rapidly undergo spontaneous WGD. From its origin within the first 50 days of the experiment, tetraploids persist for the next 950 days (nearly 5,000 generations, the current leading edge of our experiment) in ten replicate populations, despite being genomically unstable. Using synthetic reconstruction, biophysical modeling, and counter-selection experiments, we found that tetraploidy evolved because it confers immediate fitness benefits in this environment, by producing larger, longer cells that yield larger clusters. The same selective benefit also maintained tetraploidy over long evolutionary timescales, inhibiting the reversion to diploidy that is typically seen in laboratory evolution experiments. Once established, tetraploidy facilitated novel genetic routes for adaptation, playing a key role in the evolution of macroscopic multicellular size via the origin of evolutionarily conserved aneuploidy. These results provide unique empirical insights into the evolutionary dynamics and impacts of WGD, showing how it can initially arise due to its immediate adaptive benefits, be maintained by selection, and fuel long-term innovations by creating additional dimensions of heritable genetic variation.

## MAIN

Polyploidy, resulting from whole-genome duplication (WGD), is widespread in nature and is an important driver of species adaptation and diversification^1,3^. Most, if not all, living species bear signatures of ancient WGDs^3,9^. However, the establishment of nascent polyploids is rarely successful^10,11^, since newly-formed polyploids usually face fitness disadvantages when competing in a population of diploids under normal environments^5,12^. Furthermore, nascent polyploids often exhibit genomic instability^5–7^ and rapidly revert to diploidy via chromosome losses^13–15^ (but see Lu et al.^16^ for an exception to this overall trend). This costly and transient nature of nascent polyploidy raises the question of how polyploidy can rise in a diploid population and be maintained over long-term evolution. A central hypothesis is that the immediate phenotypic effects of polyploidy, often stemming from the increased size of polyploid cells, can confer fitness advantages under novel, often stressful, environments^12,17–20^. This has been suggested to contribute to many ancient WGDs that rose during periods of drastic climate change^21^. However, it remains elusive whether selection on the immediate phenotypic effects of nascent polyploidy is sufficient to drive its rise and long-term persistence, especially given the genomic instability of polyploidy that typically erodes recently-duplicated genomes.

The instability of nascent polyploid genomes may provide an evolutionary advantage under novel environments by rapidly generating genetic variation, especially via aneuploidy^9,22^. This has been shown to play an important role in the rapid evolution of microbes ^2,23–25^ and cancer ^4,26,27^. This benefit of WGD often arises as polyploids undergo genome reduction towards diploidy, rendering polyploidy a transient “gateway karyotype” towards novel, adaptive genotypes^3^. However, it remains untested whether this benefit is necessarily associated with genome reduction, or if WGD can also facilitate adaptation via novel aneuploidy when polyploidy is maintained (*i.e.*, it still possesses a baseline tetraploid genome content), potentially fueling longer-term adaptation.

Understanding how WGDs rise, are maintained, and drive both short- and long-term adaptation is fundamental to our understanding of their evolutionary impacts. Our current knowledge about WGDs is largely based on the comparison of natural polyploids with their diploid relatives, which, while informative, is confounded by evolution following WGD establishment and concomitant evolutionary processes other than WGD. Experimental evolution provides a novel opportunity to overcome these limitations. However, despite many experiments examining the evolution of synthetic diploids and polyploids (reviewed in Todd et al.^15^ and Gerstein et al.^14^), no prior work has observed how polyploids arise *de novo* in a diploid population. Moreover, the rapid losses of polyploidy in laboratory experiments limit our ability to use this approach to study the long-term maintenance and consequences of WGD. Thus, we lack an experimental system for directly observing and examining how WGDs arise *de novo* and subsequently evolve over long evolutionary timescales.

Here we report that our ongoing Multicellularity Long Term Evolution Experiment (MuLTEE)^8^ provides a unique experimental system to circumvent these long-standing constraints. Initially designed to study the evolution of a nascent multicellular organism, we subjected mixotrophic and anaerobic populations of snowflake yeast (*Saccharomyces cerevisiae*), a model of undifferentiated multicellularity, to 1000 rounds (∼5000 generations) of daily selection for larger size. Surprisingly, we found that tetraploidy rapidly evolved in our initially-diploid populations, and it has been maintained in all replicate populations for the rest of the experiment. This offers a unique opportunity to uncover the evolutionary factors underpinning the origin and maintenance of WGDs over long evolutionary timescales. We found that tetraploidy initially evolved due to its immediate phenotypic effects, generating larger clusters formed by larger, longer cells, which is adaptive under our size-based selection regime. Continuous selection for larger size also maintained tetraploidy, despite its high degree of intrinsic genomic instability, allowing it to persist for unprecedented time scales (∼4,750 generations and counting). Moreover, while mixotrophic populations remained microscopic and predominantly euploid, all anaerobic populations evolved extensive aneuploidy, which we show here played a central role in the subsequent origins of macroscopic multicellular size. Taken together, we provide direct experimental evidence that selection on the immediate phenotypic effects of polyploidy is sufficient to drive its rise and long-term maintenance, despite its intrinsic genomic instability and tendency for diploidization. Moreover, even when polyploidy is maintained, the instability inherent to polyploid genomes can also fuel long-term adaptation, via the origin and persistence of aneuploidy.

### 1000 days of multicellular evolution

We began the MuLTEE (Fig. 1a) with the goal of examining the open-ended evolution of a nascent multicellular organism^8^. The snowflake yeast ancestor we used to initiate the MuLTEE is a homozygous diploid *S. cerevisiae* with *ACE2* knockout, which causes incomplete cell separation after mitosis^28^. Snowflake yeast grow as clonal multicellular groups and reproduce through branch fragmentation induced by cell packing stress^28,29^. We evolved snowflake yeast with three metabolic treatments: mixotrophic, obligately anaerobic, and obligately aerobic, with five replicate populations (i.e., lines) per treatment. We focus on mixotrophic and anaerobic populations in this study, which are referred to as PM1-5 and PA1-5, respectively. We subjected them to daily cycles of growth and selection for rapid settling through liquid media, which select for both rapid growth and larger multicellular size. While the anaerobic snowflake yeast evolved to be ∼20,000-fold larger within 600 days (∼3000 generations), forming macroscopic clusters, the mixotrophic snowflake yeast only increased in size by ∼6-fold^8^. This was because oxygen diffusion limitation constrains the evolution of increased size under mixotrophic, but not anaerobic, metabolism^8,30^.

**Fig. 1.**
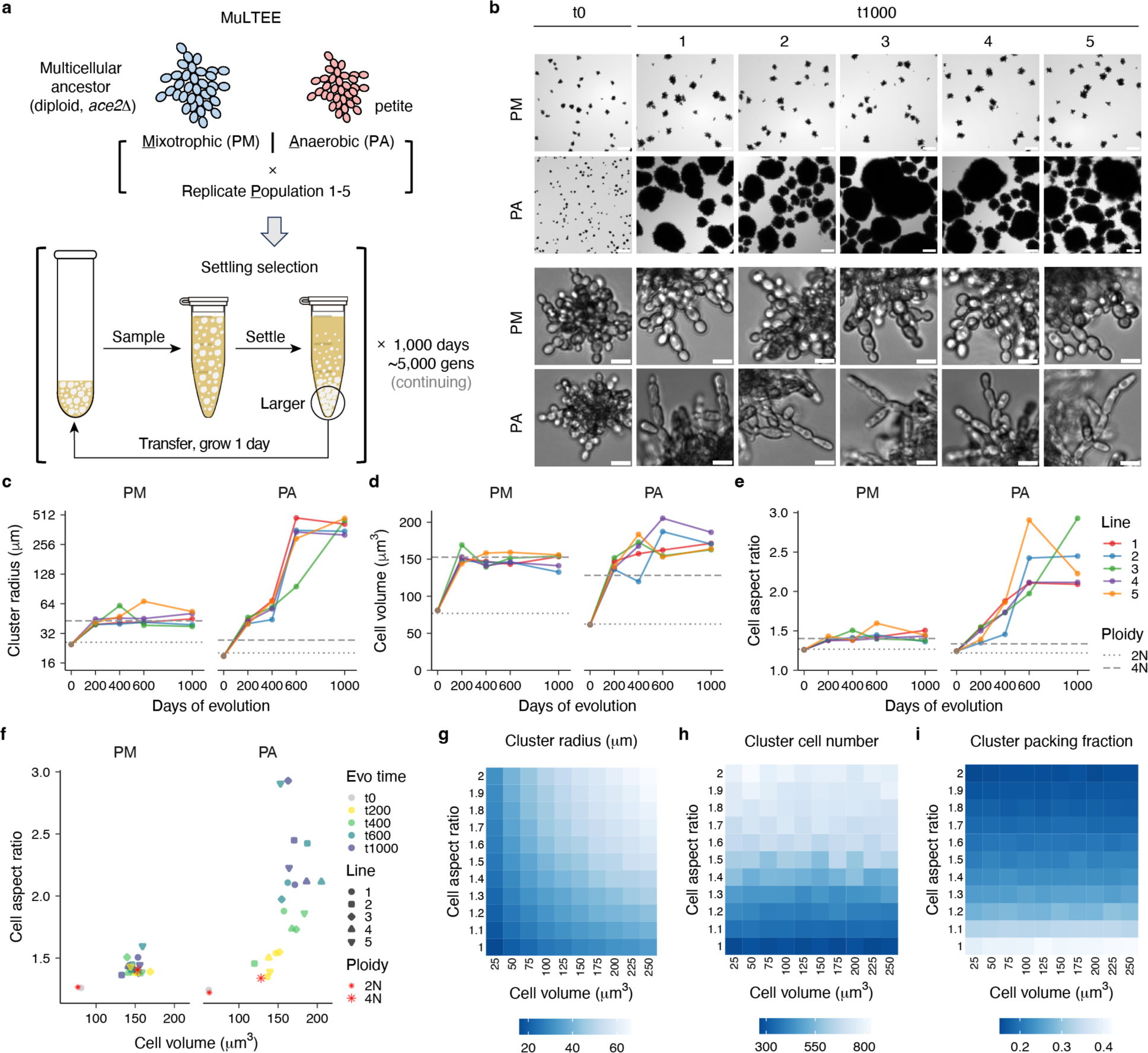
| Increased cell volume and aspect ratio drive the evolution of larger multicellular groups in the 1000-day MuLTEE. **a**, Experimental setup of the MuLTEE. **b**, Representative cluster-level (top two rows) and cell-level (bottom two rows) images of the ancestor (t0) and day-1000 (t1000) isolates from the five replicate populations evolved under mixotrophic (PM) and anaerobic (PA) conditions. Scale bars, 200 μm (cluster-level images) and 10 μm (cell-level images). **c-e**, Evolutionary dynamics of biomass-weighted mean cluster radius (Methods) (**c**), mean cell volume (**d**), and mean cell aspect ratio (**e**), showing the values of PM/PA t0 and PM/PA1-5 t200, t400, t600, and t1000 isolates (on average, n = 886 clusters (**c**) and 1288 cells (**d,e**) measured for each of the 42 strains), with the gray dotted and dashed lines representing artificially-constructed diploid and tetraploid strains (mean value of the four biological replicates in **Fig. 3c-e**), respectively. **f**, Relationship of mean cell volume and mean cell aspect ratio of strains in **c-e**. **g-i**, Heat maps showing the biophysical simulations of how cell volume and cell aspect ratio affect the mean cluster radius (**g**), cell number per cluster (**h**), and cell packing fraction (fraction of the cluster volume occupied by cells) (**i**) of clusters at fragmentation (n = 50 simulated clusters per pair of parameter values).

We have continued evolving snowflake yeast in the MuLTEE for 400 more transfers, or 1000 days (∼5000 generations) in total (Fig. 1b, Extended Data Fig. 1). In our model system, novel multicellular traits arise as an emergent property of changes in cell-level traits, with changes in cellular phenotypes (especially increased cell aspect ratio) underpinning the emergence of larger, tougher clusters^8,29–31^. Here, we characterized the evolutionary history of key group-level and cell-level traits across the first 1000 days of the MuLTEE (Fig. 1c-f, Extended Data Fig. 2). We focus particularly on the evolution of cell volume, as this has never been systematically examined in our model system and is a common phenotypic effect of WGD^9^. We found that the PMs experienced a 1.9-fold increase in cell volume within the first 200 days of our experiment (*P* = 7.53 × 10^−5^, *t*_4_ = 16.7, two-tailed one-sample *t*-test), which remained largely unchanged for the rest of the experiment (*P* = 0.637, *F*_3,16_ = 0.58, one-way analysis of variance (ANOVA)) (Fig. 1d). Similarly, the cell volume in PAs increased by 2.3-fold during the first 200 days (*P* = 1.07 × 10^−5^, *t*_4_ = 27.3, two-tailed one-sample *t*-test), with little further increase after 400 days (*P* = 0.593, *F*_2,12_ = 0.546, one-way ANOVA) (Fig. 1d). While PMs largely plateaued in cell aspect ratio and cluster size after 200 days (*P* = 0.455 and 0.613, *F*_3,16_ = 0.917 and 0.618, respectively, one-way ANOVA) following an initial increase (*P* = 1.61 × 10^−4^ and 8.27 × 10^−5^, *t*_4_ = 13.7 and 16.3, respectively, two-tailed one-sample *t*-test, comparing t200s and t0), the PAs displayed continuous increases in these two traits over the experiment (*P* = 3.17 × 10^−6^ and 5.12 × 10^−7^, *r*^2^ = 0.67 and 0.73, respectively, linear regression) (Fig. 1c,e).

Since cell volume increased concomitantly with cell aspect ratio in both PMs and early PAs (Fig. 1d-f), we sought to disentangle their effects on cluster size, using our previously-validated biophysical model^8,32^. We found that increased cell volume and aspect ratio both lead to larger clusters (Fig. 1g), but in mechanistically different ways: larger cells give rise to proportionally larger clusters without changing cell packing density or cell number per cluster, while longer cells reduce cell packing density and allow growing more cells per cluster (Fig. 1h,i). These results demonstrate how increases in cell volume and aspect ratio, two key cellular traits, contribute to the evolution of larger multicellular clusters in our snowflake yeast model system through distinct biophysical mechanisms.

### Convergent origin and maintenance of polyploidy

In most organisms, a two-fold increase in genome content increases cell volume proportionally^9^. We thus examined the ploidy of our yeast across the 1000 days of the MuLTEE. First, we sequenced the genomes of isolates from t200 and beyond, and we found that most of their point mutations have allele frequencies centered around 0.25, with the others around 0.5, 0.75, or 1 (Fig. 2a), suggesting they had evolved tetraploid genomes. To further validate this, we developed an imaging-based method for measuring ploidy levels of multicellular yeast strains (Extended Data Fig. 3a). Since asynchronous, exponential-phase cultures are used for ploidy measurements, the distribution of cellular DNA contents of a single, clonal strain contains two peaks, corresponding to G1- and G2-phase cells (Extended Data Fig. 3b). Using this method, we confirmed that all the evolved isolates indeed have near-tetraploid DNA contents, while their ancestors are diploids (Fig. 2b). To trace the origin of tetraploidy, we measured ploidy distributions of populations from earlier time points, and we discovered that in all ten populations, tetraploidy had emerged and become dominant as early as 50 days and had become fixed by 100 days (Fig. 2c). Together, these results show that in all ten PM and PA lines, tetraploidy emerged very early in the MuLTEE, and it has been maintained for the subsequent 950 transfers (and counting). While the PM lines were mainly euploid, the PA lines evolved extensive aneuploidy (Fig. 2d, Extended Data Fig. 4), which we will examine in detail later in the paper.

**Fig. 2.**
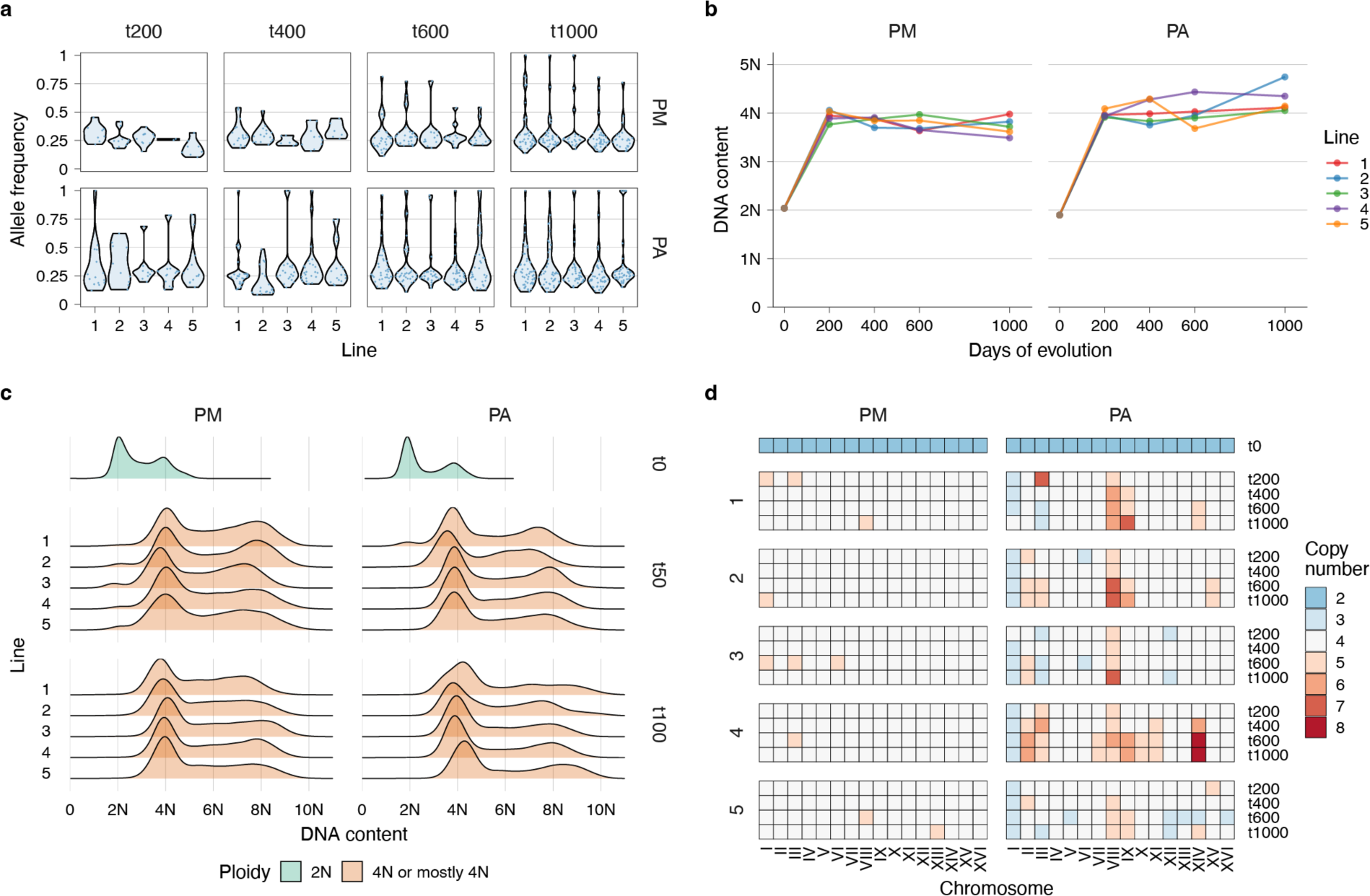
| Tetraploidy rapidly emerged and was maintained in all PMs and PAs; PAs subsequently evolved extensive aneuploidy. **a**, Violin plot of the distribution of allele frequencies (each determined as the fraction of the read number of the mutant allele in the read number of all alleles) of novel point mutations identified in PM/PA1-5 t200, t400, t600, and t1000 isolates, with each dot representing one point mutation. **b**, Evolutionary dynamics of DNA content, showing that of PM/PA t0 and the strains used in **a** (on average, n = 11456 cells measured for each of the 42 strains). **c**, Ridge plot of the distribution of cellular DNA contents in the PM/PA t0 and PM/PA1-5 t50 and t100 populations (on average, n = 12922 cells measured for each of the 22 populations). Since asynchronous cultures were used for ploidy measurements, G1- and G2-phase cells of a single genotype in a population could form two distinct DNA-content peaks. **d**, Heat maps of the karyotypes of PM/PA t0 and the strains used in **a**, determined based on chromosome coverages from the whole-genome sequencing data.

To our knowledge, this is the first evolution experiment in which polyploidy convergently evolved from diploid ancestors. Importantly, polyploidy is not directly induced by our experimental conditions, as in some prior experiments^33,34^. Moreover, the long-term persistence of polyploidy in our experiment (>4750 generations) is in dramatic contrast to previous evolution experiments, where, under various conditions, tetraploid yeast ancestors are genomically unstable and typically converge to diploidy within a few hundred generations^2,13,16,35,36^. This raises the question: what drove the origin and maintenance of polyploidy in the MuLTEE?

### Polyploidy confers an immediate selective advantage to snowflake yeast

In unicellular *S. cerevisiae*, increased ploidy is known to cause increases in both cell volume^6,37^ and cell aspect ratio^37^. In fact, increased cell size is a universal feature of polyploid cells across eukaryotes^9^. As increased cell volume and aspect ratio both contribute to larger snowflake yeast clusters (Fig. 1g), we hypothesize that tetraploidy arose in the MuLTEE because it brings immediate phenotypic effects, generating larger, longer cells that yield larger clusters, which is beneficial under settling selection. To test this directly, we genetically engineered tetraploidy in a diploid snowflake yeast background (Extended Data Fig. 5a). Consistent with our hypothesis, under both mixotrophic and anaerobic conditions, tetraploid clusters consist of larger, more elongated cells and are larger than their diploid counterparts (Fig. 3a-e, Extended Data Fig. 5b-e). Interestingly, the scaling relationship between increased cell volume and increased cell aspect ratio due to tetraploidization is nearly identical across mixotrophic and anaerobic conditions (slopes = 0.0018 µm^−3^ in both cases) (Fig. 3f), suggesting that similar mechanisms may underlie how tetraploidy impacts cellular morphology under both metabolic conditions.

**Fig. 3.**
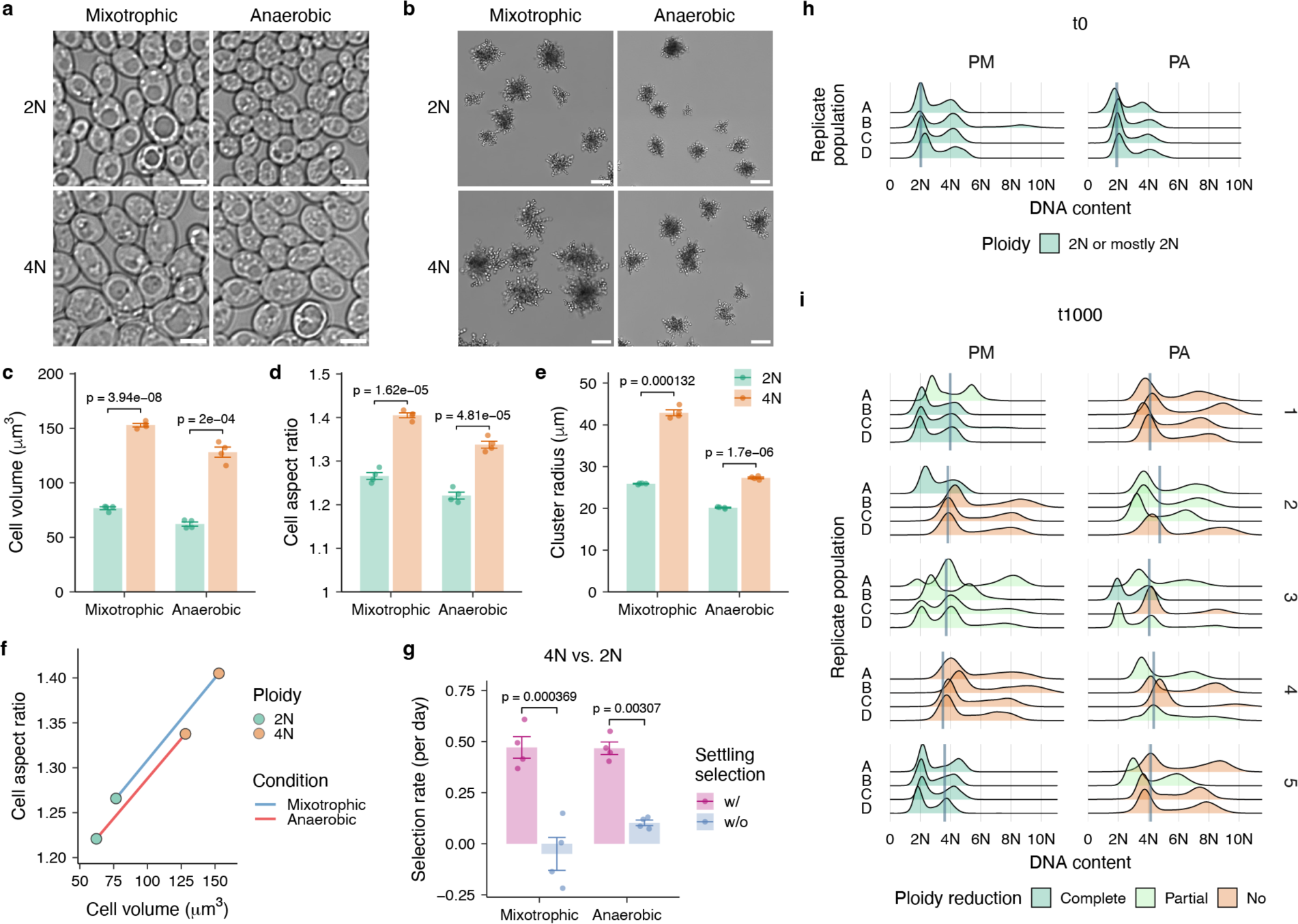
| Tetraploidy confers immediate phenotypic effects that are beneficial under selection for larger size, driving the origin and maintenance of tetraploidy despite its genomic instability. **a,b**, Cell-level (**a**) and cluster-level (**b**) images of the engineered diploid and tetraploid clusters under mixotrophic and anaerobic conditions. Scale bars, 5 μm (**a**) and 50 μm (**b**). **c-e**, Mean cell volume (**c**), mean cell aspect ratio (**d**), and biomass-weighted mean cluster radius (**e**) of the strains in **a,b**. **f**, Relationship of the changes in cell volume and cell aspect ratio due to tetraploidization, under mixotrophic and anaerobic conditions. Values are means of the four replicates or petite mutants in **c,d**. **g**, Per day selection rate of competing engineered tetraploid clusters against diploid counterparts with and without settling selection for three days, under mixotrophic and anaerobic conditions. For **c-e,g**, values are mean ± s.e.m. (n = 4 biological replicates for mixotrophic condition or 4 independent petite mutants for anaerobic condition; on average, 2458 cells (**c,d**) and 1077 clusters (**e**) were measured per replicate or petite mutant, and 381 clusters (**g**) were measured per sample). *P* values were calculated by two-tailed Welch’s t-test. **h,i**, Distribution of cellular DNA contents in the evolved populations, initiated with PM/PA t0 (**h**) and PM/PA1-5 t1000 isolates (**i**) with four replicate populations (A-D), after selecting against larger size for 70 days (on average, n = 14487 cells measured per population). Gray vertical line, ancestral DNA content of each strain from **Fig. 2b** (corresponding to G1 peak). Levels of ploidy reduction: “complete”, reduction to diploidy; “partial”, reduction to an intermediate ploidy level between diploidy and tetraploidy, or a mixture of tetraploidy and lower ploidy levels; “no”, no detectable ploidy reduction.

Next, we compared the engineered tetraploid strains with the evolved isolates to see how much of the phenotypic changes in the MuLTEE can be explained by tetraploidization alone. In PMs, engineered tetraploidy recapitulates the increases in cluster size, cell volume and cell aspect ratio in the MuLTEE (Fig. 1c-e). Even over the full 1000 days, there appears to be little phenotypic adaptation in this treatment beyond what is conferred by tetraploidy. In PAs, tetraploidy contributed to most of the cell volume increase in the first 200 days of the experiment (Fig. 1d), as well as most of the increases in cluster size and cell aspect ratio in the first 50 days (Extended Data Fig. 5f,g), when tetraploidy became dominant (Fig. 2c). However, tetraploidy alone does not explain the subsequent increases in cluster size and cell aspect ratio in PAs (Fig. 1c,e).

To examine whether tetraploidy indeed confers an immediate fitness benefit under selection for larger size, we competed engineered diploid and tetraploid clusters for three days with and without settling selection, using a label-free fitness assay (Extended Data Fig. 6). Supporting our hypothesis, under both mixotrophic and anaerobic conditions, settling selection strongly favored tetraploidy (Fig. 3g), increasing its average frequency from 53% to 82% within three days.

### Polyploidy arises and persists due to sustained selection for larger size

Our results above suggest that tetraploidy evolved as a mechanism of increasing group size. To test this more directly, we re-evolved the MuLTEE ancestors with selection acting in the opposite direction. Specifically, we grew and transferred our yeast on solid media (Extended Data Fig. 7a), which prior work demonstrated strongly favors smaller groups due to lower within-group competition for resources^38^. We evolved four replicate populations of each of PM and PA ancestors on agar with daily dilution for 70 days (∼500 generations). All eight populations remained predominantly diploid (Fig. 3h). This is markedly different from the MuLTEE, in which tetraploid strains became dominant within the first 50 days (∼250 generations) of settling selection (Fig. 2c). This also indicates that tetraploidization is not an inherent tendency of our model system and does not evolve without appropriate selection.

While tetraploidy initially evolved as a mechanism to increase group size in our experiment, nascent tetraploidy, especially in *S. cerevisiae*, is notoriously unstable^2,13,16,35,36^. What explains its maintenance over long-term evolution (nearly 5000 generations) in the MuLTEE? We have two central hypotheses: tetraploidy may be actively maintained by selection for larger size, or, alternatively, tetraploid yeast could have evolved molecular mechanisms that stabilize the duplicated genomes, like in Lu et al^16^.

To disentangle these hypotheses, we performed a similar reverse selection experiment, evolving the tetraploid PM and PA t1000 isolates on solid media where larger size is selected against. We evolved four replicate populations of each of the ten t1000 isolates on agar with daily transfers for 70 days (∼500 generations) (Extended Data Fig. 7a). In total, 22/40 evolved populations exhibited ploidy reduction after 500 generations of counter-selection, including nine populations that completely reverted to diploidy (Fig. 3i). This time scale is similar to previous evolution experiments that observed rapid ploidy reduction of tetraploid unicellular yeast under various conditions^2,13,16,35,36^. Further, ploidy reductions were observed in 8/10 genetically-distinct t1000 isolates. This suggests that tetraploid snowflake yeast remained genomically unstable throughout the MuLTEE, and that the maintenance of a baseline tetraploid genome throughout the experiment was due to sustained selection for larger multicellular size. Notably, not all the 20 populations initiated with PM t1000 isolates reduced in ploidy or cluster size within 70 days, but reductions in ploidy and in cluster size were always correlated, and all populations that completely diploidized showed mean cluster sizes that reduced to the level of diploid PM ancestor (Extended Data Fig. 7b).

### Aneuploidy underpins the origin of macroscopic size

Polyploidy is associated with much higher rates of aneuploidy than diploidy^39–41^. Nascent polyploids tend to rapidly undergo genome reduction, and the resulting aneuploidy can offer a novel genetic route for rapid adaptation to novel environments^2,25^. However, it remains untested whether polyploids, when maintained, can also adapt by generating aneuploidy. The MuLTEE, with its long-term maintenance of polyploidy, presents a unique system to directly test this hypothesis. Consistent with the instability of tetraploid genomes, we observed cases of aneuploidy (Fig. 2d) and segmental aneuploidy (Extended Data Fig. 4) in the evolved isolates in the MuLTEE. Notably, while all PM isolates were euploid or close to euploid, all PA isolates displayed extensive aneuploidy (Fig. 2d). This distinction is correlated with the divergent multicellular evolution under the two metabolic treatments: while PMs remained microscopic over 1000 days, PAs underwent sustained multicellular adaptation, evolving remarkably larger multicellular groups (Fig. 1c). We thus hypothesize that the evolved aneuploidy in PAs might have contributed to their origin of macroscopic multicellular size, arguably the most striking phenotypic innovation that evolved in the MuLTEE so far.

Parallel changes in aneuploidy and multicellular size in PAs over the MuLTEE suggest a potential link between the two. The level of aneuploidy in PAs generally increased over evolution (linear regression against time for number of aneuploid chromosomes, total number of chromosome copies deviating from euploid tetraploidy, and coefficient of variation of chromosome copy numbers, *P* = 0.023, 0.017, and 0.020, *r*^2^ = 0.10, 0.11, and 0.11, respectively) (Fig. 2d), mirroring the trends in cluster size and cell aspect ratio (Fig. 1c,e). However, after the evolution of macroscopic size (which occurred by t600 for all lines except PA3), these complex karyotypes stayed remarkably conserved between t600 and t1000 in each line (Fig. 2d). The exception, PA5, where considerably different karyotypes were observed between its t600 and t1000 isolates (Fig. 2d), also displayed substantially different cluster sizes and cell aspect ratios between them (Fig. 1c,e). These observations suggest that aneuploidy might have played a critical role in the evolution of macroscopic size in PAs.

To test this hypothesis more directly, we first attempted to genetically reconstruct specific aneuploidy using the conditional centromere method (p*GAL1*-*CEN*)^42,43^. However, this method exhibited low efficiency and off-target chromosome changes in our system. Therefore, we turned to a forward-genetics approach, selecting for losses of macroscopic size in evolved isolates and examining if karyotype changes are involved. This is enabled by the accidental finding that the heritable change in colony morphology on agar plates is a strong indicator of spontaneous losses of macroscopic size: all macroscopic PA isolates typically form ring-shaped colonies (“donut” colonies), interspersed with rare, larger, flattened colonies (“spread” colonies) that can no longer form macroscopic clusters when cultured in liquid media (Fig. 4a). Such donut-to-spread transitions happened rapidly, as spread colonies could be readily observed after plating an overnight culture grown from a donut colony. Previous studies reported that rapid changes in yeast colony morphology can be the result of karyotype changes^44^. Thus, to systematically examine if karyotype changes are involved in losses of macroscopic size, we randomly selected three donut colonies from each of the nine macroscopic PA t600 and t1000 isolates (excluding the still microscopic PA3 t600 isolate) and then isolated one spread colony derived from each donut colony. In total, we prepared 25 donut-spread pairs, including only two pairs from PA1 t600 isolate (because the donut strain in the third pair did not grow macroscopic in later experiments) and PA5 t1000 isolate (because we were only able to obtain two pairs after many attempts). All spread strains had dramatically reduced cluster size, no longer forming macroscopic clusters (Fig. 4b). Consistent with the previously demonstrated effects of cell-level traits on cluster size, the donut-to-spread transitions were associated with overall decreases in cell volume, and more significantly and consistently, in cell aspect ratio (Fig. 4c-f).

**Fig. 4.**
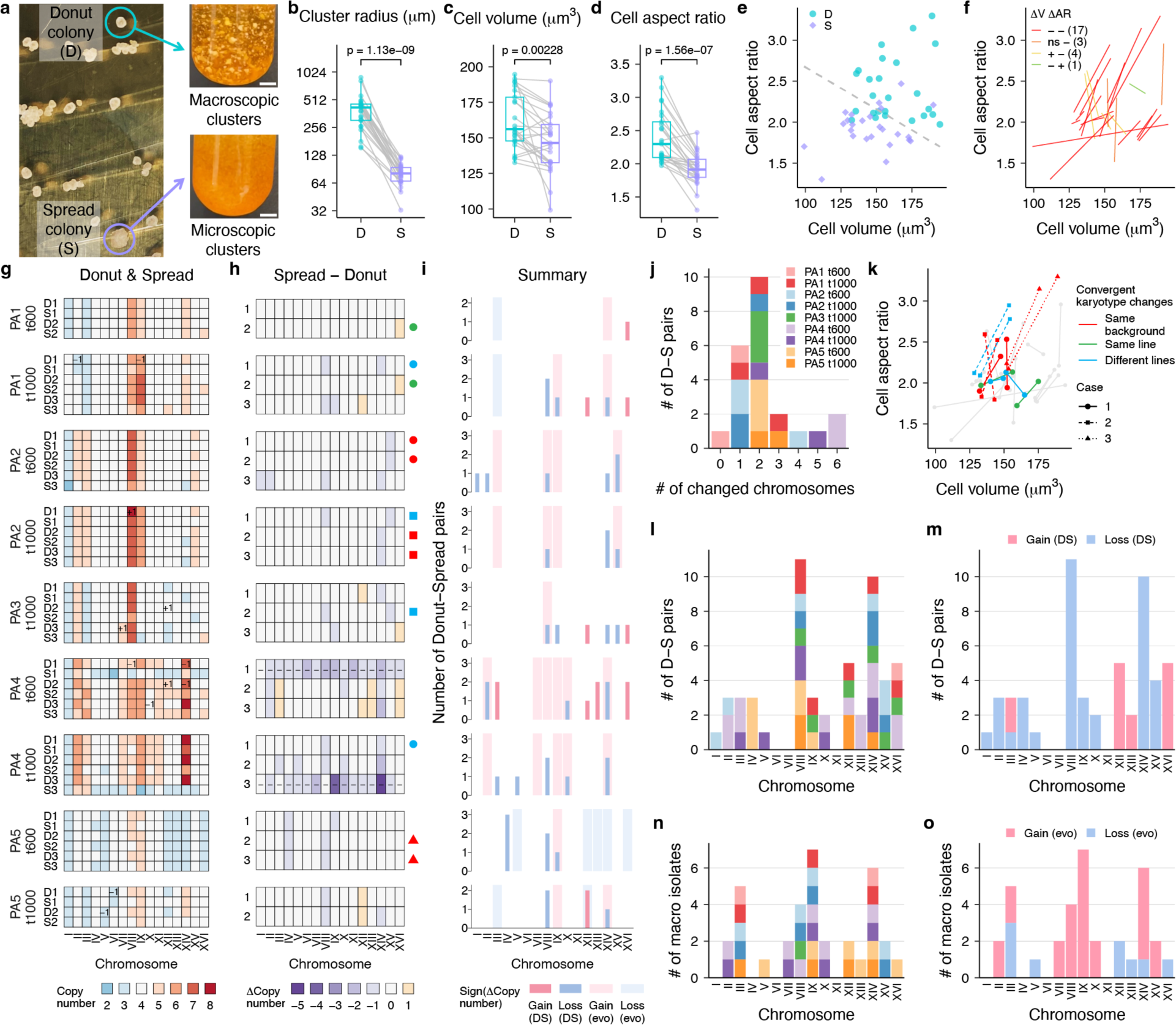
| Aneuploidy promotes the origin of macroscopic multicellularity in PAs. **a**, Evolved macroscopic isolates (PA3 t1000 is shown) form “donut” colonies (D) and occasionally “spread” colonies (S) that cannot form macroscopic clusters. Scale bar, 5 mm. **b-d**, Biomass-weighted mean cluster radius (**b**), mean cell volume (**c**), and mean cell aspect ratio (**d**) of the 25 D-S pairs (each connected by a line) from nine macroscopic isolate backgrounds. Boxes, interquartile range (IQR); center lines, median; whiskers, values within 1.5 × IQR of the first and third quartiles. *P* values were calculated by two-tailed paired t-test (on average, 605 clusters (**b**) and 1294 cells (**c,d**) measured per strain). **e,f**, Relationship of how cell volume and aspect ratio changed in D-S transitions, with donut and spread strains separated by a linear boundary generated by support vector machine (**e**), and with each D-S transition connected by a line and grouped into four classes based on the signs of changes in cell volume and cell aspect ratio (*P* < 0.05, two-tailed Welch’s t-test with Benjamini-Hochberg correction; ns, not significant; D-S pair counts in each class are indicated in brackets) (**f**). **g,h**, Karyotypes of D-S pairs (**g**) and karyotype changes in D-S transitions (**h**). Inside the heatmaps, numbers (**g**) show chromosome copy number differences between donut strains and their backgrounds in **Fig. 2d**, and dashes (**h**) indicate two D-S transitions with near-triploidization that were excluded from subsequent analyses. Symbols beside the heatmap (**h**) indicate cases of convergent karyotype changes where two D-S pairs came from the same background (red), different backgrounds but the same PA line (green), or different lines (blue), with each case in a different symbol shape. **i**, For each background, chromosomes with copy number changes in the evolution (light rectangular backgrounds) and loss (dark bars, showing number of occurrences) of macroscopic size. **j**, Histogram of the number of chromosomes with copy number changes in each D-S pair, colored by background. **k**, Changes in cell volume and aspect ratio compared between D-S pairs with convergent karyotype changes, with symbol color-shape code consistent with **h** and line color-shape code similarly assigned. **l-o**, Aggregating all backgrounds in **i**, number of times each chromosome changes its copy number in the loss (**l,m**) and evolution (**n,o**) of macroscopic size, colored by background (**l,n**, share color legend with **j**) or direction of change (**m,o**).

To identify the genetic changes underlying the spontaneous losses of macroscopic size, we sequenced and compared the genomes of each donut-spread pair. We found no strong statistical evidence that point mutation changes played a systematic role in this process (Extended Data Fig. 8, Supplementary File 1). In contrast, 24 of 25 donut-to-spread transitions were associated with karyotype changes (Fig. 4g,h). Two spread strains in PA4 even lost almost an entire set of chromosomes and partially triploidized (Fig. 4g,h), which may explain why they possess the smallest cell volumes among all spread strains (Fig. 4c), given the effect of ploidy level on yeast cell volume^6,37^. We excluded these two extreme cases from subsequent analyses, due to their divergence from our otherwise tetraploid background and confounding effect of dramatically decreased DNA content. Karyotype changes in spread strains were mostly limited to copy number changes in 1-2 chromosomes (Fig. 4j), and among the 53 chromosome copy number changes in total, all but two changed by only one copy (Fig. 4h).

Surprisingly, despite the large number of ways that karyotypes could potentially change, we observed six cases of convergent karyotype changes among just 23 donut-spread pairs (*P* = 8.24 × 10^−6^, probability of observing six or more cases with identical karyotype changes, each with changes in no more than two chromosomes, assuming equal probability of gains and losses of each of the 16 chromosomes) (Fig. 4h). Three of these cases of convergent changes even involved donut-spread pairs derived from different genetic backgrounds (either different evolutionary time points in the same line or different lines) (Fig. 4h). Most of these convergent karyotype changes were also associated with similar changes in cell volume and aspect ratio (Fig. 4k), suggesting a link between karyotype changes and phenotypic changes. Moreover, we also observed convergent changes in the copy numbers of certain chromosomes across donut-to-spread transitions (Fig. 4l), and strikingly, each chromosome (except for chromosome III) that underwent copy number changes always changed in the same direction across all spread strains (i.e., a chromosome was either always gained or always lost) (Fig. 4m). This is most clearly reflected in chromosome VIII and XIV, which were convergently lost in all five lines and in almost half of all spread strains (*P* = 6.28 × 10^−7^ and 5.05 × 10^−6^, respectively, binomial probability of observing 11 or more, or 10 or more losses of any given chromosome among 53 chromosome copy number changes, assuming equal probability of gains and losses of each of the 16 chromosomes) (Fig. 4l,m). Taken together, the prevalent and convergent changes at the karyotype and chromosome level during donut-to-spread transitions, as well as their correlations with phenotypic changes, strongly suggest that changes in aneuploidy underpin the spontaneous losses of macroscopic size in the donut-to-spread transitions.

To investigate the role of aneuploidy in the evolution of macroscopic size in the MuLTEE, we compared the chromosome copy number changes associated with the emergence of macroscopic size during the MuLTEE to those involved in the spontaneous loss of macroscopic size in our donut-to-spread colony transitions. We first identified the candidate chromosomes that are directly involved in the evolution of macroscopic size in each line, by identifying the chromosomes with copy number changes that are absent in any of the microscopic isolates (t200 and t400, and for PA3, also t600) but present in at least one of the macroscopic isolates (t600 and t1000). This yields one set of candidate chromosomes for each of the nine macroscopic isolates (Fig. 4i, light background). Notably, for 13 of 18 candidate chromosomes (exceptions mostly in PA5), the evolutionary changes of their copy numbers observed in the t600 isolate were maintained in the t1000 isolate of the same line (Fig. 4i). Among the 53 chromosome copy number changes in all donut-to-spread transitions, 20 occurred in the candidate chromosomes of the corresponding isolate background. Of these, all changed in the opposite direction to their evolutionary changes in the MuLTEE (Fig. 4i, contrasting colors in dark bar and light background), suggesting that copy number changes of these candidate chromosomes contributed to the evolution of macroscopic size. Importantly, these changes of candidate chromosomes in donut-to-spread transitions cannot be solely explained by them occurring at a high rate and hitchhiking with other mutations, otherwise we would expect them to change in the same direction over the MuLTEE, while we observed exactly the opposite, implying that their changes over evolution were the result of selection, not mutational bias.

Furthermore, when comparing across the five PA lines, we found that some candidate chromosomes and their directions of copy number changes over evolution are shared among multiple lines (e.g., chromosome VIII, IX, XIV), while the other candidate chromosomes are more line-specific (Fig. 4n,o). Strikingly, in the donut-to-spread transitions, all the chromosomes (candidate or not) that changed copy numbers in more than one spread strain changed consistently in the opposite direction to their evolutionary changes (comparing Fig. 4m and Fig. 4o). Moreover, as the copy numbers of some candidate chromosomes can rapidly change in the donut-to-spread transitions, their maintenance from t600 to t1000 isolates in the MuLTEE suggest that, just like polyploidy, at least some part of the evolved aneuploidy was maintained by continuous selection for larger multicellular size despite its intrinsic instability.

## Discussion

The Multicellularity Long Term Evolution Experiment (MuLTEE) was initially designed to study the open-ended evolution of increasingly complex multicellular life. Surprisingly, our results reveal that the MuLTEE also serves as a unique polyploidy long-term evolution experiment, providing direct empirical insights into the short- and long-term evolutionary dynamics of nascent whole-genome duplications (WGDs). We found that tetraploidy evolved in our system due to its immediate phenotypic effects, producing larger, longer cells that give rise to larger multicellular clusters, which is adaptive under our selection regime. Despite its intrinsic genomic instability, tetraploidy was maintained over thousands of generations by daily selection for larger organismal size. However, when this selection pressure was removed, even highly-evolved tetraploids rapidly reverted towards diploidy, highlighting the critical role that environmental selection can play in stabilizing nascent WGDs. Further, we discovered that extensive, convergently-evolving, and evolutionarily-conserved aneuploidy played a key role in the subsequent evolution of macroscopic multicellularity in our model system.

Our results have several important implications for understanding the dynamics of WGD. First, we show that nascent polyploidy is not necessarily transient, even prior to the evolution of genome-stabilizing mechanisms. WGD is long known to bring immediate phenotypic changes, many stemming from increased cell size (a universal feature of polyploid cells and central to evolved polyploid isolates in the MuLTEE), which can confer fitness benefits under novel environments^12,17,18,20,21^. Here, we experimentally demonstrate that selection favoring the immediate phenotypic effects of WGD is sufficient to drive both its origin and maintenance, despite the intrinsic instability of polyploid genomes. As a result, the duration of the environment favoring polyploidy may be a key determinant of whether a nascent WGD will be transient or persistent. This may also explain the increased abundance of polyploids in certain environments^14,45^ (e.g., tetraploid yeast in bakeries^46^). Importantly, unlike cases where genome-stabilizing mechanisms evolve, the polyploid genomes actively maintained under environmental selection can still be intrinsically unstable, rapidly reverting towards diploidy in the absence of the polyploidy-favoring selection. This is consistent with the fact that most ancient WGDs eventually reverted to diploidy^3^.

Building upon previous works showing that WGD can profoundly influence evolutionary dynamics by altering the rate, spectrum and fitness effect of mutations^2,14,15,25^, our experiment is the first to directly examine how recently duplicated genomes evolve when polyploidy is maintained over long evolutionary timescales. In the MuLTEE, we find that the point mutations arising in the polyploid background are highly heterozygous, and polyploidy, even when maintained, can still exhibit genomic instability and generate aneuploidy, exploring novel karyotypic space and facilitating long-term adaptation. Moreover, our findings challenge the prevailing notion that aneuploidy serves as a temporary solution to novel environmental conditions^39,47^. Instead, we demonstrate that even complex aneuploid karyotypes can be maintained under sustained selection pressures for thousands of generations, despite the inherent instability of aneuploid genomes. This observation also offers an alternative explanation for the aneuploidy frequently found in natural polyploids^39^: rather than being a transient, non-adaptive consequence of genomic instability, in some cases they may be the result of selection, preserved over long timescales when the selective environment remains stable.

Phylogenetic evidence shows that ancient WGDs often precede periods of increased species diversification, albeit with a time delay^48^. Our work provides a potential ecological explanation for this phenomenon: in novel environments, selection may favor and maintain polyploids, allowing structural variation and highly heterozygous point mutations to accrue. When the environment no longer favors polyploidy, relatively rapid diploidization can lead to the dramatic rearrangement of these accumulated mutations, generating vast genetic diversity that may fuel adaptive radiation.

While previous work has revealed the adaptive roles of aneuploidy mostly in promoting stress resistance and virulence^39,40^, our results show that aneuploidy can also contribute to the evolution of morphological innovations (*i.e.*, macroscopic multicellularity), expanding its functional repertoire. Further work will be required to explore the mechanistic basis of how aneuploidy drives this phenotypic change. Possible mechanisms may include gene dosage effects, where changes in chromosome copy number alter the expression levels of key genes involved in cell morphology and multicellular development, and epistatic interactions between structural variations and point mutations. From an evolutionary perspective, it will be interesting to examine whether the divergent karyotypes evolved and maintained in different PA lines result from historical contingency and represent local fitness optima in the karyotypic space. Additionally, it is worth investigating why the aneuploidy in PMs is limited. PMs do not appear to be constrained, however, in their intrinsic capacity of generating aneuploidy, as most PM t1000 isolates underwent rapid ploidy reduction when evolving on agar (Fig. 3i). The near-euploidy in PMs may instead be because the evolution of increased multicellular size is constrained under this condition^8,30^, and aneuploidy can incur deleterious effects^49^ and may not evolve if its benefits do not outweigh its costs.

The MuLTEE is now the longest-running evolution experiment not just in the evolution of multicellularity, but also in the evolution of polyploidy, offering direct empirical insights into how WGDs can rise, become maintained, and drive short- and long-term adaptation. We anticipate that this open-ended, multi-condition, long-term evolution experiment will continue to serve as a source of inspiration and a testbed for various hypotheses regarding WGD, one of the most fundamental processes in eukaryotic evolution.

## METHODS

### Multicellularity Long Term Evolution Experiment (MuLTEE)

The details of how we constructed the ancestor strains and conducted experimental evolution with settling selection up to 600 days were described previously^8^. Briefly, we constructed a multicellular diploid *S. cerevisiae* strain from a unicellular diploid Y55 strain by *ACE2* deletion. From this grande strain which grows both aerobically and anaerobically in YPD media (1% yeast extract, 2% peptone, 2% dextrose), we isolated a spontaneous petite mutant which grows anaerobically in YPD media despite the presence of oxygen. We evolved five replicate populations of mixotrophic and anaerobic snowflake yeast (referred to as PM1-5 and PA1-5, respectively) with daily selection for increased size over 1000 days. Every day, we grew each population in 10 mL of YPD media at 30°C with shaking at 250 rpm for 24 hours, and then we transferred 1.5 mL of the culture into a 1.5 mL Eppendorf tube, let clusters settle on the bench for 3 minutes and transferred the bottom 50 µL into 10 mL of fresh YPD media for the next day of growth. Once PA1-5 evolved macroscopic clusters that settle rapidly, we used wide-bore pipette tips to minimize breaking clusters during pipetting, and we also decreased the settling time to 30 seconds to allow sustained selection for increased size, with this change occurring on ∼350 days for PA2 and 5 and ∼500 days for PA1, 3, and 4. For simplicity, starting from day 850, we sampled 1 mL, instead of 1.5 mL, from each culture for daily settling selection. We archived a glycerol stock for each population every 10-15 days. We extracted one representative clonal isolate from each of the ten populations archived on day 200, 400, 600, and 1000 for subsequent analyses.

### Measuring cluster size

We revived strains from glycerol stocks by growing them on YPD plates at 30°C for 2 days. Then we inoculated each strain into 10 mL of YPD media and grew them at 30°C with 250 rpm shaking for three days with daily settling selection before transferring to fresh media, recapitulating how the strains grow during the evolution experiment. On the last day, after settling selection and transfer, we grew the cultures for 24 hours and sampled them at 4 hours (exponential phase) and 24 hours (stationary phase) for measuring cluster size. Unless otherwise noted, the 24-hour measurements are used throughout the paper, as they represent the states of the cultures right before settling selection.

Prior to imaging, we gently shook each culture by hand (without vortexing, which may break clusters) and added an appropriate volume of the culture (1-250 µL, depending on cluster density) into H_2_O containing 10 µL of 16% (w/v) formaldehyde (Thermo Scientific, 28906) in a 24-well plate, making up 510 µL per well. For macroscopic strains, we sampled the culture using manually-cut wide-bore 1 mL tips to avoid breaking macroscopic clusters during pipetting, and we typically sampled twice from each culture and transferred into two wells to reduce the randomness in sampling macroscopic clusters. We gently shook the 24-well plate to evenly spread out the clusters and allowed 5-10 minutes for clusters to settle down.

We used a Nikon Eclipse Ti inverted microscope to scan the whole wells by taking and stitching 5 × 8 brightfield, shading-corrected images with 10% overlap at 4x magnification, scanning at 1 mm/second. We developed a semi-automated image analysis pipeline using ImageJ v1.54f to (1) segment clusters using auto local thresholding (which allows detecting clusters ranging from tiny branches to macroscopic clusters) and split touching clusters using seed-based watershed, (2) perform manual correction using a custom, user-friendly toolkit for improving the speed and reproducibility of manual correction, and (3) measure the cross-sectional area of each cluster. We removed cluster objects with an area below 40 µm^2^. Using R v4.1.2, we converted cluster areas to cluster volumes and radiuses, treating clusters as perfect spheres. For each strain, we calculated its biomass-weighted mean cluster radius^8^ by first calculating its mean cluster volume weighted by cluster volume and then converting it to radius.

### Measuring cell volume and aspect ratio

We revived strains from glycerol stocks by growing them on YPD plates at 30°C for 2 days. Then we inoculated each strain into 10 mL of YPD media and grew them at 30°C with 250 rpm shaking for three days with daily settling selection before transferring to fresh media. On the last day, we transferred 100 µL of culture (without settling selection) to 10 mL of fresh media and grew it for 12 hours, following Bozdag et al.^8,30^.

To prepare samples for imaging, we transferred 25 µL of each culture to a 1.5 mL microcentrifuge tube. For macroscopic strains, we sampled the culture using 100 µL wide-bore tips to allow pipetting macroscopic clusters, and we broke clusters into tiny pieces using 100 µL regular-bore tips to facilitate crushing them into a single cell layer later. We pelleted the clusters by spinning at 5000 × g for 1 minute, washed them with 1 mL of 1× PBS, and incubated them in 100 µL of 5 µM Calcofluor White (a blue fluorescent cell wall stain) in 1× PBS at room temperature. We gently crushed 5 µL of stained clusters (without shearing, which can lyse cells) into a single cell layer between a microscope slide and a coverslip.

We used a Nikon Eclipse Ti inverted microscope to take brightfield and fluorescent (UV channel) images of at least five fields of view (FOVs) per strain at 40x magnification. For each FOV, we set the focus based on the brightfield channel by manually moving to a z-plane where cells look gray (rather than brighter or darker) and then moving down by 0.7 µm to get sharp fluorescent signals of the cell wall. We developed a semi-automated image analysis pipeline using Cellpose v2.2.2^50^ and ImageJ v1.54f to perform (1) automated cell segmentation using Cellpose (specify cell diameter as 80 pixels for images at 0.073 µm/pixel), run on GPUs provided by Georgia Tech’s Phoenix Cluster, (2) manual correction using Cellpose’s Graphical User Interface (GUI), and (3) measurement of the area, lengths of major and minor axis, aspect ratio, and solidity of each cell in ImageJ. We removed cell objects with an area below 1 µm^2^ or a solidity below 0.8. Using R v4.1.2, we calculated cell volumes using the formula V = 4/3 × πab^2^ (treating yeast cells as perfect ellipsoids), where a and b are the lengths of the major and minor axis of each cell, respectively.

### Biophysical simulation

We adapted our previously published biophysical model^8,32^ to disentangle and elucidate the effects of cell volume and cell aspect ratio on cluster size. In our model, we added new cells (modeled as prolate ellipsoids) in the characteristic snowflake yeast budding pattern until a certain total amount of overlap between cells is reached (representing the threshold for cluster fragmentation), whereupon the simulation is terminated. Using this model, we swept through a range of overlap thresholds to find a value that recapitulated our empirical data. Then, using that value, we swept through a range of cell volumes (from 25 to 250 µm^3^) and aspect ratios (from 1 to 2). We simulated 50 clusters at each pair of parameter values, and measured cluster volume (estimated by the volume of the convex hull bounding the cluster), cell number, and packing fraction of each simulated cluster upon fragmentation.

### Whole-genome sequencing and analysis

To sequence yeast genomes, we first extracted yeast genomic DNA using DNA purification kit (VWR). Genomic library preparation using Illumina DNA Prep kit and sequencing using Illumina NovaSeq 6000 and NovaSeq X Plus sequencer with 150-bp paired-end reads were performed by the Microbial Sequencing Center at University of Pittsburgh. We obtained FASTQ files of genome reads at an average coverage of 160X.

To analyze the genome reads, we first filtered and trimmed low-quality reads using Trimmomatic v0.39. Next, we aligned the reads to the S288C reference genome^51^ (R64-3-1 build) using BWA-MEM v0.7.17^52^. Following GATK’s Best Practices pipeline^53^, we generated binary alignment files (BAM) and marked duplicates using Picard Toolkit v2.27.5. Next, we used GATK Haplotypecaller v4.2.4.1 to call for variants. To filter out variants with low genotype quality, we used VCFTOOLS v0.1.16^54^. Next, we extracted novel mutations that are only present in the evolved genomes by comparing them against the ancestral genome via bcftools-isec v1.10^55^. Finally, we annotated novel mutations using snpEff v5.0^56^ (including assigning mutation impacts as high, moderate, low, or modifier) and calculated their allele frequencies by dividing the read depth of the mutant allele with the total read depth in each particular position.

To determine the karyotype of each strain, we first computed the read depth of each base position (base coverage) in the genome using Bamtools Stats v2.5.1 and then estimated chromosome copy numbers in R v4.1.2. Specifically, we calculated the mean base coverage in each 1kb non-overlapping bin along each chromosome (bin coverage), and then normalized the bin coverages in each genome by dividing it with the median bin coverage in the whole genome. Next, we calculated bin copy number by multiplying the normalized bin coverage with the baseline ploidy level of the strain (determined by imaging-based ploidy measurement below). Finally, we estimated the copy number of each chromosome as its rounded median bin copy number.

For donut and spread strains, we also estimated mutation allele copy number in R v4.1.2, using mutation allele frequency and chromosome copy number calculated above. For a mutation with its corresponding chromosome having copy number N, its allele copy number would be 0 or N if its allele frequency is 0 or 1, respectively. For other values of allele frequency, we estimated the mutation allele copy number (C) as the integer between 1 and N-1 where C/N is the closest to the allele frequency. In cases of ties (observed in 7 out of 2003 mutations in 46 donut and spread strains), we chose the allele copy number that has the least change from the other strain in the corresponding donut-spread pair. We also used C/N as the corrected allele frequency.

To calculate the distribution of mutation impacts of a point mutation randomly introduced into the yeast genome (S288C reference genome, R64-3-1 build), we only considered single-nucleotide substitutions, as multi-nucleotide substitutions and indels are much rarer and were not observed in the mutations that were gained in donut-to-spread transitions. We calculated the probability of the random mutation having high, moderate, low, or modifier impact as the probability of it being a nonsense, missense, synonymous or intronic, or non-coding/intronic mutation.

### Yeast strain construction

To construct isogenic grande diploid and tetraploid snowflake yeast, we started with two *S. cerevisiae* grande haploid unicellular strains (*MAT*a *hoΔ*::*hphNT1* and *MAT*α *hoΔ*::*hphNT1*), derived from Y55 background (from which the MuLTEE ancestors were also derived). We first deleted the *ACE2* open reading frame from these two haploid strains using kanMX cassette, which made them multicellular, and then mated them to form a grande diploid strain (a/α). Next, we transformed this diploid strain with the p*GAL1*-*HO*-*natNT2* plasmid containing *CEN/ARS* elements, and then transiently induced *HO* expression by galactose to switch the mating type from a/α to a/a and α/α (determined by halo assay), followed by selection for plasmid loss. We then mated the a/a and α/α diploid strains to form a grande tetraploid strain (a/a/α/α).

To generate petite diploid and tetraploid snowflake yeast, we isolated spontaneous petite mutants from the grande diploid and tetraploid strains, respectively, by selecting colonies that are smaller than normal colonies on a YPD plate (1% yeast extract, 2% peptone, 2% dextrose, 1.5% agar) and cannot grow on a YP-Glycerol plate (the same as YPD plate but with the dextrose replaced by 2.5% glycerol). Since petite mutation is not isogenic, we isolated four independent petite mutants per strain, each of which is derived from a different single colony of the parental grande strain.

### Imaging-based ploidy measurement of snowflake yeast

Common protocols for ploidy measurement of unicellular yeast are based on flow cytometry of DNA-stained single cells, where fluorescent intensity of the DNA stain like propidium iodide (PI) scales linearly with DNA content^15,57^. However, these protocols cannot be readily applied to multicellular yeast without an efficient method for dissociating clusters into single cells, so we developed an imaging-based protocol for measuring the ploidy level of snowflake yeast strains. Basically, we used fluorescent images of PI-stained, flattened clusters to quantify the PI intensity of G1-phase nuclei, which is compared to strains with known ploidy to estimate the DNA content of the focal strain.

We adapted the sample preparation procedure prior to fluorescent imaging from Todd et al^57^. Importantly, in every experiment, two control strains with known ploidy, namely, the engineered grande diploid and tetraploid snowflake yeast, also went through the same procedure of sample preparation and imaging. We first grew each strain of interest in YPD media to mid-log phase, and then transferred 250 µL of the culture into a 1.5 mL microcentrifuge tube. We pelleted the clusters by spinning at 5000 × g for 1 minute and washed them with 1 mL of H_2_O. For macroscopic strains, we broke the clusters into small pieces by pipetting vigorously using 100 µL regular-bore tips before washing. We fixed and permeabilized the cells in 1 mL of 70% ethanol at room temperature for 2 hours with end-to-end rotation on a mini-rotator (BioSan, Bio RS-24) at the maximum speed, followed by washing with 1 mL of 50 mM sodium citrate twice. We resuspended the clusters in 200 µL of 50 mM sodium citrate containing 0.5 mg/mL RNase A (MP Biomedicals, 101076) and incubated them in a 37°C heat block for 2 hours with gentle inversion every 30 minutes (as clusters settle over time). After RNA digestion, we added 5 µL of 1 mg/mL PI (Thermo Fisher, P1304MP) and incubated the mixture in a 30°C incubator in dark overnight with rotation on a mini-rotator (BioSan, Bio RS-24) at the minimum speed to keep clusters from settling. PI-stained clusters can be stored at 4°C for no longer than a week before imaging.

For fluorescent imaging, we crushed 5 µL of PI-stained clusters into a single cell layer between a microscope slide and a coverslip. For each sample, we took 14-bit images of ∼10 FOVs at 20x magnification using a Nikon Eclipse Ti inverted microscope. For each FOV, we set the focus by manually moving to a z-plane where cells look gray (rather than brighter or darker) in the brightfield channel and then moving down by 1 µm to get sharp fluorescent signals of the nuclei. We imaged the flattened clusters in the red fluorescent channel (exposure 600ms, gain 2.2x) at the focal plane as well as one z-plane 0.3 µm above and below the focal plane (three z-planes in total) to detect the PI-stained nuclei, and then imaged in the brightfield channel (exposure 100ms, gain 4.1x) at the focal plane. Importantly, we set the exposure and gain of the fluorescent channel such that the brightest pixels in the tetraploid control strain are ∼80% of the maximal allowed pixel value while minimizing photobleaching and noise.

We performed quantitative image analysis using ImageJ v1.54f. We first performed maximum intensity projection of the three z-planes taken in the fluorescent channel and used the resulting image for segmentation and fluorescence quantification. We segmented nuclei and filtered them to include only single round nuclei. We measured the total PI fluorescence intensity of each nucleus with background subtraction, where the background fluorescence is the median PI fluorescence intensity of the cytoplasm in the cluster the focal nucleus is in. The resulting nuclear PI intensity scales linearly with the DNA content.

We analyzed the image analysis results in R v4.1.2. For each sample, we removed tiny objects with areas two median absolute deviations below median (nucleus segmentation artifacts), and we manually removed FOVs with outlier distributions of nuclear PI intensity. The distribution of nuclear PI intensity in a clonal strain contains two peaks that correspond to G1- and G2-phase cells. We estimated the DNA content of each nucleus by dividing its PI intensity with the PI intensity of a haploid genome, which was estimated by averaging across the two ploidy control strains, i.e. mean(G1 peak intensity of diploid control strain / 2, G1 peak intensity of tetraploid control strain / 4). We estimated the DNA content of a clonal strain as the DNA content of its G1 peak.

### Competition assay

To perform competition assays between the engineered diploid and tetraploid snowflake yeast without fluorescent labeling (which may incur fitness cost), we developed an imaging-based method for distinguishing clusters with different ploidy levels, utilizing the fact that tetraploid clusters contain larger cells than diploid counterparts. Specifically, prior to imaging, we transferred an appropriate volume (12-18 µL, depending on cluster density) of the culture containing diploid and tetraploid clusters into a 24-well plate with 500 µL of H_2_O per well. We gently shook the 24-well plate to evenly spread out the clusters and allowed 5-10 minutes for clusters to settle down. To image the center of each well, we used a Nikon Eclipse Ti inverted microscope to take and stitch 6 × 6 brightfield, shading-corrected images with 5% overlap at 20x magnification, scanning at 1 mm/second. We imaged at three z-planes, including the z-plane where the cells touching the well bottom look slightly bright, as well as 5µm and 10µm above it. Using ImageJ v1.53q, we first segmented the clusters using the middle z-plane (with manual correction), and then we segmented the bright cells in the cluster edges in all three z-planes, thus capturing cells in different focal planes. Next, for each cluster, we calculated the mean area of the five largest cells detected across all three z-planes. This value is sufficient to distinguish the engineered diploid and tetraploid clusters in either grande or petite form, with the cutoff determined in a preliminary test.

To compete the engineered diploid and tetraploid snowflake yeast under mixotrophic and anaerobic conditions, we competed grande diploid versus grande tetraploid with four biological replicates, and we paired the four diploid and tetraploid petite mutants to form four competing pairs as our four replicates. We first revived the strains from glycerol stocks by growing them on YPD plates at 30°C for 2 days. Then we inoculated four replicate tubes for each grande strain (diploid and tetraploid) as well as one replicate tube for each petite mutant (diploid and tetraploid), and we grew them in 10 mL of YPD media at 30°C with 250 rpm shaking for 2 days with daily 1:100 dilution. To start the competition of each of the eight competing pairs, we mixed 500 µL of the 24-hour culture of each competing strain (1 mL in total), from which we inoculated 100 µL into 10 mL of YPD media in each of two culture tubes and grew them for three days, transferring daily with and without settling selection, respectively. For competitions with settling selection, we recapitulated the settling selection scheme in the MuLTEE: every day we transferred 1.5 mL of each 24-hour culture into a 1.5 mL microcentrifuge tube, let clusters settle for 3 minutes, and transferred the bottom 50 µL into 10 mL of YPD media to grow for another day. For competitions without settling selection, we grew each culture with daily 1:100 dilution. We measured the frequency of the total cluster volume in each competing strain at the start and the end of the 3-day competition, using the above imaging-based method to distinguish diploid and tetraploid clusters. Finally, we calculated the per-day selection rate of tetraploid versus diploid strain, using the formula r = [log(% of tetraploids on day 3 / % of tetraploids on day 0) - log(% of diploids on day 3 / % of diploids on day 0)] / 3 days.

### Evolution experiment with selection against larger size

We revived PM/PA t0 and PM/PA1-5 t1000 isolates from glycerol stocks by growing them on YPD plates at 30°C for 2 days. Then we inoculated each of the 12 strains into 10 mL of YPD media and grew them at 30°C with 250 rpm shaking overnight, from which we transferred 20 µL to inoculate each of the four replicate populations for the evolution experiment. We grew the 48 populations in a 30°C static incubator for 70 days (∼500 generations) in a spatially-structured environment using 24-well plates containing 2 mL of YPD agar per well. Every 24 hours, we resuspended each population in 1 mL of saline solution (0.85% NaCl), pre-diluted it in a microcentrifuge tube, and then transferred 20 µL of the diluted resuspension onto fresh YPD agar and spread it out by gently rocking the 24-well plate. We initially used a pre-dilution factor of 1:2 (for a total dilution of 1:100 after accounting for the volume plated), but we increased it to 1:4 (total 1:200) on day 36 after the populations had adapted to this selection regime. We prepared glycerol stocks of each population every 7 days by mixing 800 µL of the undiluted saline resuspension with 400 µL of 70% glycerol and stored them at −80°C. Prior to routine protocols of measuring ploidy level and cluster size, we revived the populations by scraping a big chunk from the glycerol stocks and growing them in 10 mL of YPD to saturation for 2 days (without transferring after the first 24 hours).

### Isolating donut and spread colonies

To isolate one pair of donut and spread colonies from a macroscopic strain, we first streaked the strain from glycerol stocks onto a YPD plate and grew it at 30°C for 2 days. Importantly, as streaking or plating a multicellular strain does not ensure that a colony is founded by a single cell, we used chitinase (Sigma, C8241) to break clusters down to single cells (though some tiny branches exist after digestion, they should have gone through a recent single-cell bottleneck) before plating cells and isolating donut or spread colonies below.

To isolate a donut colony, we picked a single donut colony on the YPD plate and transferred it into a 1.5 mL microcentrifuge tube containing 800 µL of chitinase solution (0.25 mg/mL chitinase in 50 mM potassium phosphate buffer, pH 6.0). We broke apart the colony by pipetting with 100 µL regular-bore tips and vortexing, and then incubated it at 30°C for 8 hours with gentle rotation on a mini-rotator (BioSan, Bio RS-24) at level 5. After chitinase digestion, we washed the cells twice with 1 mL of H_2_O and spread them on YPD plates with three dilution factors (1:400, 1:1000, and 1:2500). We incubated the plates at 30°C for 3 days. Then we picked a single donut colony from the plates and grew it in 10 mL of YPD media at 30°C with 250 rpm shaking for 24 hours, from which we prepared glycerol stocks by mixing 500 µL of culture and 500 µL of 70% glycerol. This served as our donut isolate.

To isolate a spread colony, we transferred 100 µL of the above 24-hour culture into a 1.5 mL microcentrifuge tube, washed it twice with 1 mL of H_2_O and resuspended it in 800 µL of chitinase solution. We performed chitinase digestion, washing, plating, and plate incubation in the same way as above for the donut isolate, except for plating at dilution factors of 1:800, 1:2000, and 1:5000 with three plates each. From these nine plates that contained ∼1000 colonies, we could usually find more than one spread colony. However, since chitinase does not perfectly digest clusters into single cells, it is challenging to accurately estimate the frequency of donut-to-spread transitions. We picked one single spread colony and streaked it on a YPD plate to confirm the spread colony morphology and to apply an additional round of clonal isolation. We incubated the plate at 30°C for 2 days. Then we picked a single spread colony, and we grew it in YPD media and prepared glycerol stocks in the same way as above for the donut isolate. This served as our spread isolate for the corresponding donut isolate. Importantly, since each spread colony was isolated from plating the 24-hour liquid culture inoculated from a donut colony, we limited the number of cell generations between donut and spread phenotypes, which allowed us to better pinpoint the relevant genetic changes and showed how rapidly macroscopic sizes can be lost.

### Statistics

We performed statistical tests using R v4.1.2, with the details described in the main text and figure legends. We implemented support vector machine in Fig. 4e using scikit-learn v1.3.2 in Python v3.11.6. We ran the biophysical simulations in MATLAB R2019a.

## Data availability

Underlying data used to make figures and raw data are available at GitHub (https://github.com/ktong25/WGD_in_MuLTEE). Raw Illumina sequencing reads are available at the NIH Sequence Read Archive under accession number PRJNA943273 (for MuLTEE evolved isolates) and PRJNA1093477 (for donut and spread strains). Raw microscopy images are archived in Ratcliff Lab’s Dropbox and are available on request.

## Code availability

Code used for image analysis and biophysical modeling are available at GitHub (https://github.com/ktong25/WGD_in_MuLTEE). Code used for conventional genome sequence analysis, data analysis, statistical tests and figure generation are archived in Ratcliff Lab’s Dropbox and are available on request.

## Acknowledgements

We thank the Microbial Sequencing Center at University of Pittsburgh for sequencing the genomes. We thank the Partnership for an Advanced Computing Environment (PACE) at the Georgia Institute of Technology for providing the research cyberinfrastructure resources and services. We thank all members of the Ratcliff Lab for feedback on the study. This work was supported by NIH grant R35-GM138030 to W.C.R. and Human Frontiers Science Program grant RGY0080/2020 to W.C.R.

## Author contributions

K.T. and W.C.R. conceived of the project. K.T., W.C.R., P.L.C. and S.D. designed the experiments. G.O.B. performed the Multicellularity Long Term Evolution Experiment, analyzed point mutations in the evolved isolates, and identified the genomic signature of tetraploidy. V.C., D.J.H. and K.T. genetically engineered tetraploid snowflake yeast. S.G., H.L.Y. and P.L.C. performed the evolution experiments with selection against larger size and measured the evolved populations. S.D. isolated donut and spread strains, characterized their phenotypes, and analyzed their point mutations. T.C.D. performed biophysical simulations. D.T.L. performed genomic DNA extraction and helped with genome analysis. K.T. performed the rest of the experiments, analyzed the data, and made the figures. K.T. prepared the first draft of the paper, and all the authors contributed to the revision.

## Supplementary information

Supplementary File 1: Analysis of point mutation changes in donut-to-spread transitions.

Supplementary Table 1: A list of strains, primers and plasmids used in the study.

## Extended Data Figures

**Extended Data Fig. 1.**
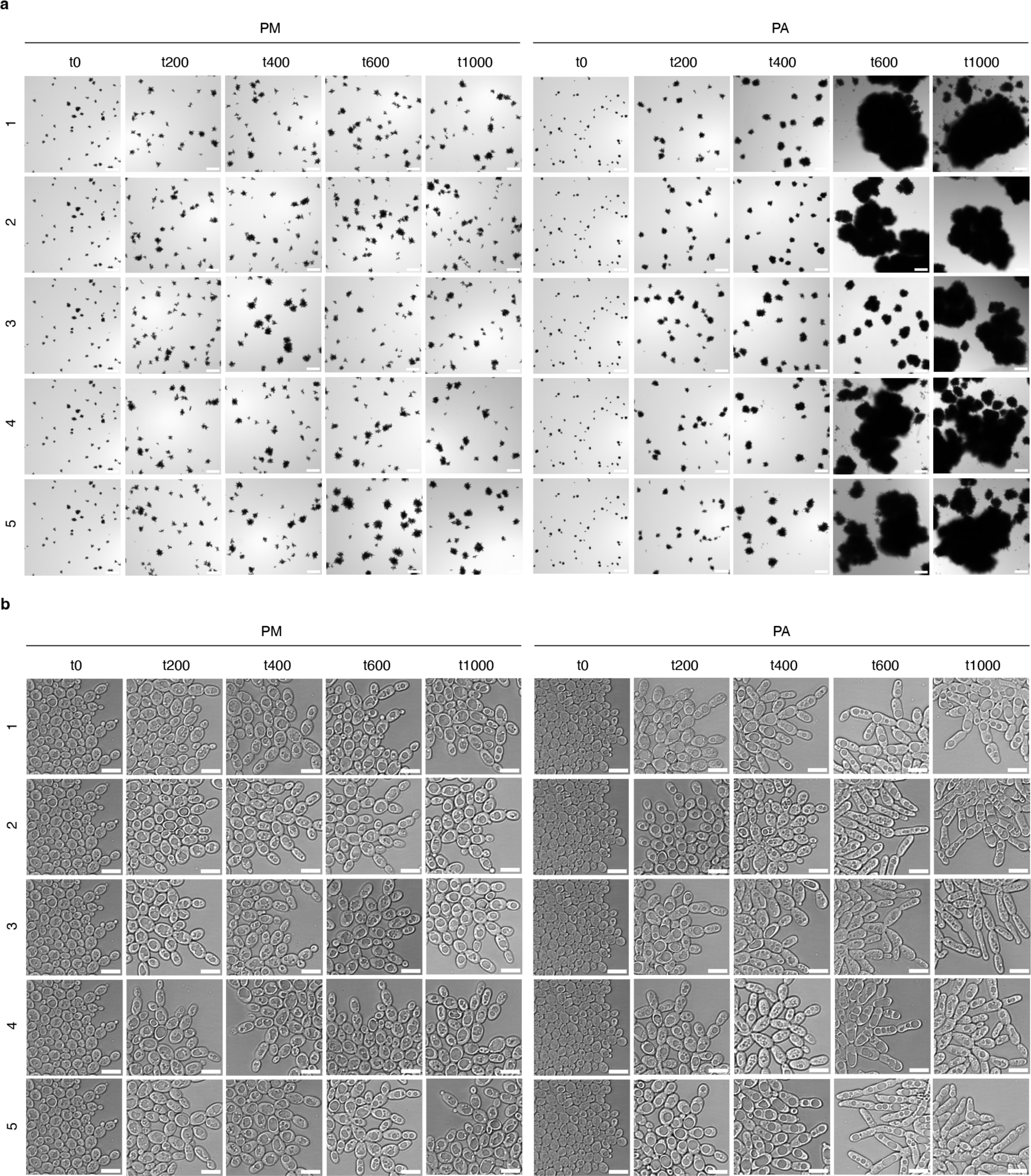
| Images of the ancestors and evolved isolates in the MuLTEE. **a,b**, Representative cluster-level (**a**) and cell-level (**b**) images of PM/PA t0 and PM/PA1-5 t200, t400, t600, and t1000 isolates. The images of PM/PA t0 are reused for five replicate populations. Scale bars, 200 μm (**a**) and 10 μm (**b**).

**Extended Data Fig. 2.**
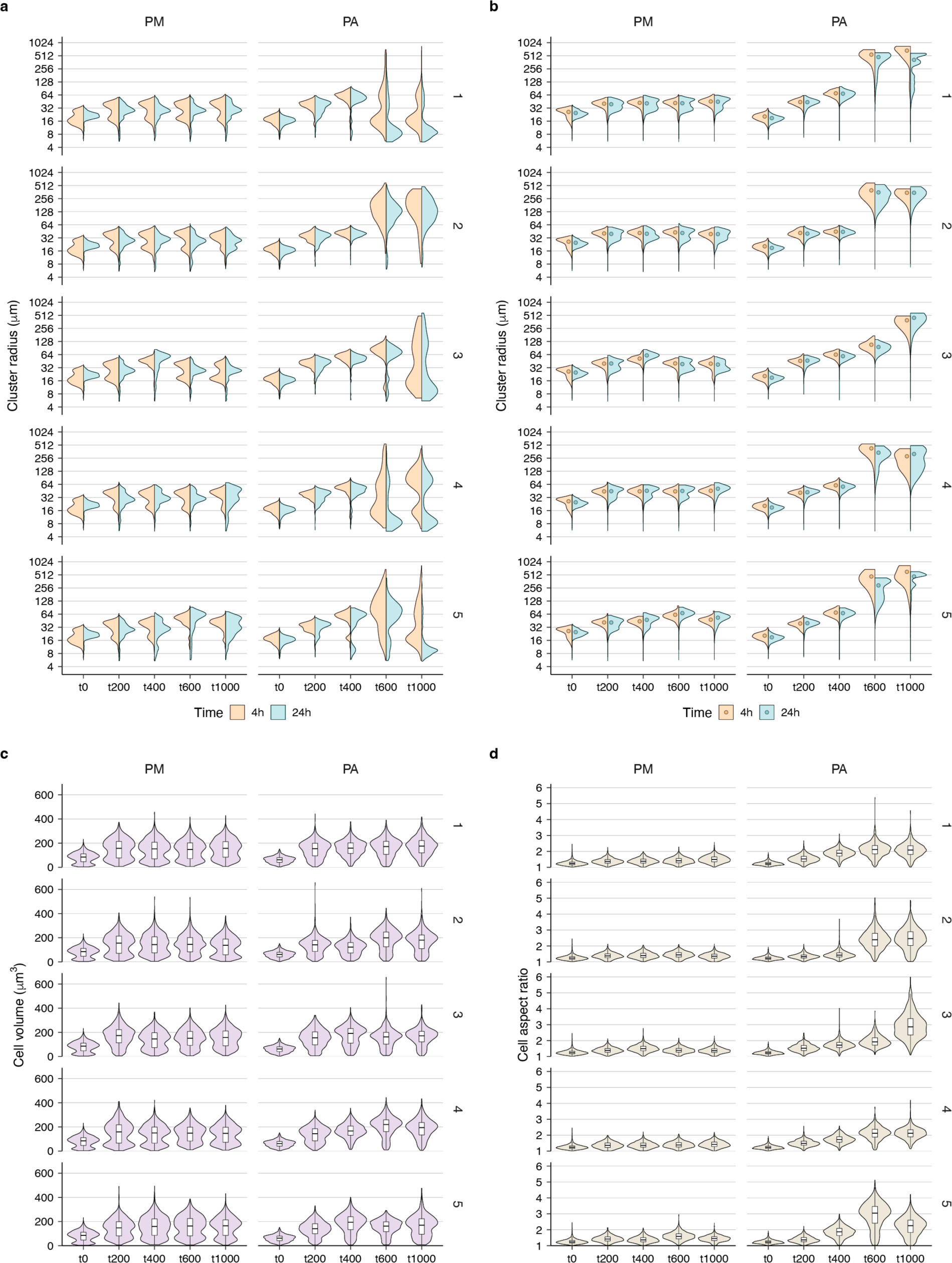
| Phenotypic characterization of the ancestors and evolved isolates in the MuLTEE. **a-d**, Violin plots showing the distributions of cluster radius (**a,b**, where **b** is weighted by cluster volume), cell volume (**c**), and cell aspect ratio (**d**) in PM/PA t0 and PM/PA1-5 t200, t400, t600, and t1000 isolates (on average, n = 789 clusters (**a,b**) and 1288 cells (**c,d**) measured per sample). The distributions of PM/PA t0 are reused for five replicate populations. For **a,b**, we measured cluster radius at 4 hours (exponential phase) and 24 hours (stationary phase) after transferring the culture to fresh media, and the 24-hour measurements (corresponding to the state of the cultures right before settling selection) are used throughout the paper unless otherwise noted. For **b**, filled circles show biomass-weighted mean cluster radius (the 24-hour values are the same as the values in **Fig. 1c**). For **c,d**, boxes, IQR; center lines, median; whiskers, values within 1.5 × IQR of the first and third quartiles.

**Extended Data Fig. 3.**
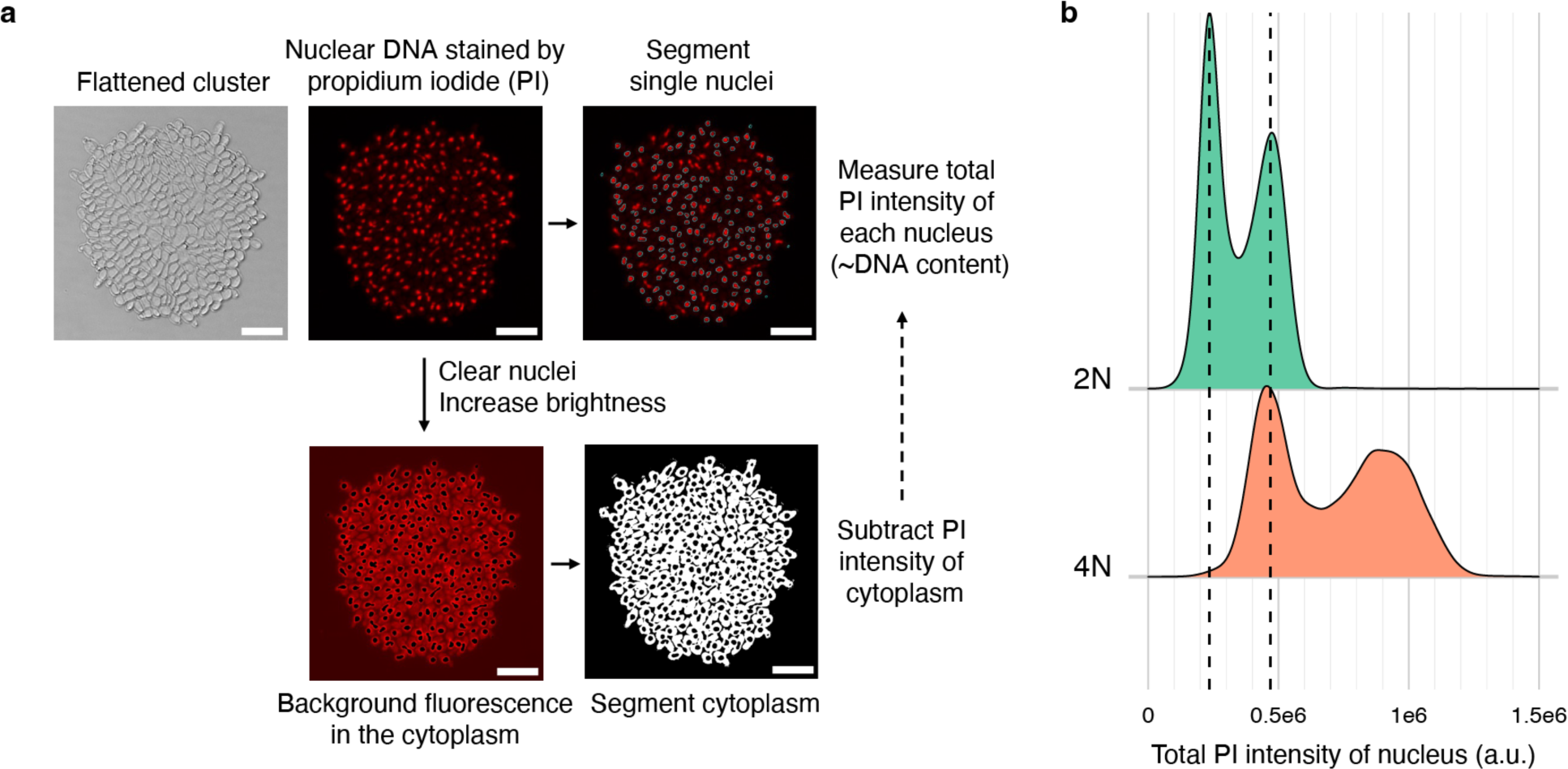
| Imaging-based method for measuring ploidy level of snowflake yeast. **a**, Overview of imaging and image analysis workflow. Snowflake yeast clusters are crushed into a single cell layer and imaged at the brightfield channel and fluorescent channel, with the latter showing the nuclear DNA stained by propidium iodide (PI). The nuclei in the fluorescent image are segmented and filtered to get single round nuclei, outlined in cyan. The fluorescent image is also nuclei-cleared and brightness/contrast-enhanced to show the background fluorescence in the cytoplasm, and the cytoplasm is segmented, shown in white, for background subtraction. The total fluorescence intensity of PI in each nucleus is quantified and background-subtracted. Scale bar, 20 µm. **b**, Distribution of the nuclear PI intensity (arbitrary unit) of the engineered diploid and tetraploid mixotrophic clusters (n = 14276 and 10031 nuclei, respectively), as a validation for this ploidy measurement method. Since asynchronous, exponential-phase cultures are used for ploidy measurements, each strain shows two peaks that correspond to G1- and G2-phase nuclei of the actively-dividing cells, and the G2 peak has double of the fluorescent intensity of the G1 peak. Also, the G2 peak of diploid clusters aligns nicely with the G1 peak of tetraploid clusters.

**Extended Data Fig. 4.**
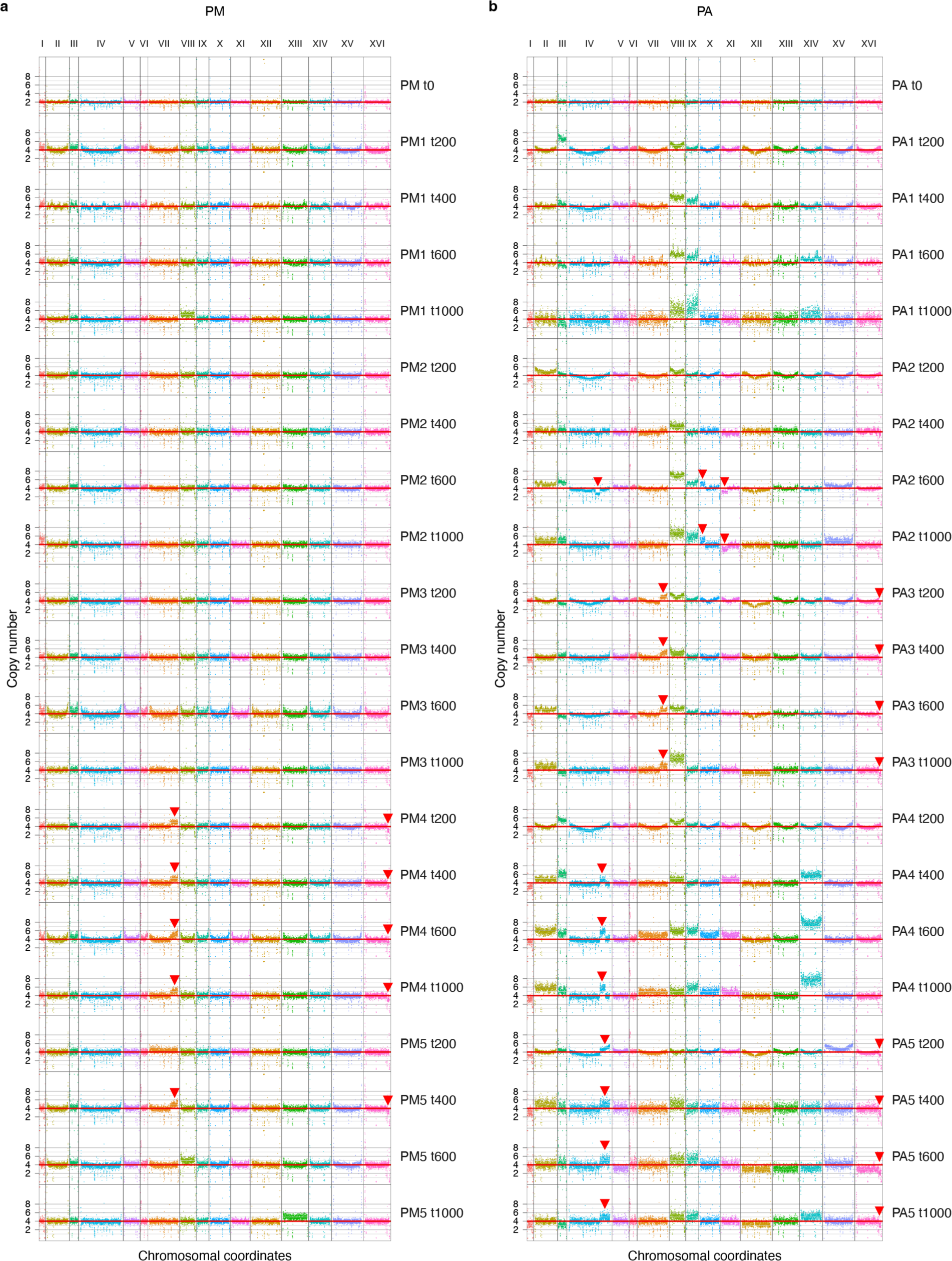
| Copy number variation of the ancestors and evolved isolates in the MuLTEE. **a,b**, Estimated copy number of each 1kb non-overlapping bin in each chromosome in PM/PA t0 and PM/PA1-5 t200, t400, t600, and t1000 isolates (**a**, PM; **b**, PA). Estimated bin copy numbers above 12 are shown as 12, indicated by little triangles. Red horizontal line, baseline ploidy of each strain (i.e., 2 for PM/PA t0 and 4 for all evolved isolates). Red arrowhead, incidence of segmental aneuploidy.

**Extended Data Fig. 5.**
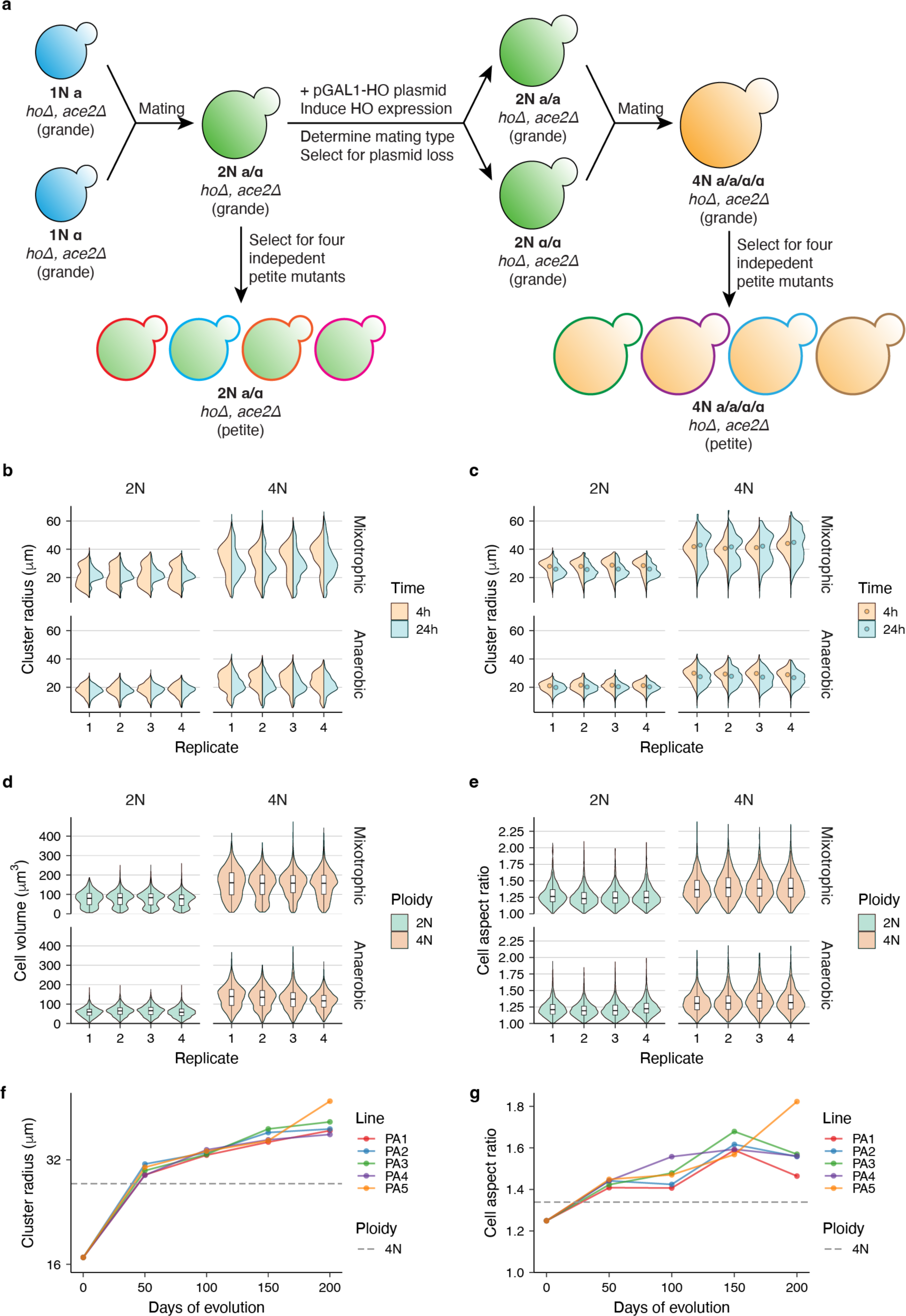
| Genetic construction and phenotypic characterization of diploid and tetraploid clusters. **a**, Procedure for engineering isogenic grande diploid and tetraploid clusters, from each of which four independent petite mutants were isolated. Isolating multiple petite mutants is important because petite mutations are not isogenic and may confound ploidy-phenotype map. Grande and petite clusters correspond to mixotrophic and anaerobic conditions, respectively. **b-e**, Violin plots showing the distributions of cluster radius (**b,c**, where **c** is weighted by cluster volume), cell volume (**d**), and cell aspect ratio (**e**) in engineered diploid and tetraploid clusters under mixotrophic and anaerobic conditions (on average, n = 922 clusters (**b,c**) and 2458 cells (**d,e**) measured per sample). Four biological replicates were measured for the mixotrophic condition, and the four independent petite mutants (each with one biological replicate) were measured for the anaerobic condition. For **b,c**, we measured cluster radius at 4 hours (exponential phase) and 24 hours (stationary phase) after transferring the culture to fresh media, and the 24-hour measurements are used throughout the paper unless otherwise noted. For **c**, filled circles show biomass-weighted mean cluster radius (the 24-hour values are the same as the values in **Fig. 3e**). For **d,e**, boxes, IQR; center lines, median; whiskers, values within 1.5 × IQR of the first and third quartiles. **f,g**, Comparison of the biomass-weighted mean cluster radius (**f**) and mean cell aspect ratio (**g**) of the engineered petite tetraploid clusters (mean of the four independent petite mutants, the same values as those in **Fig. 1c,e**) to the PA t0 and PA1-5 t50, t100, t150, and t200 populations (data from Bozdag et al. 2023).

**Extended Data Fig. 6.**
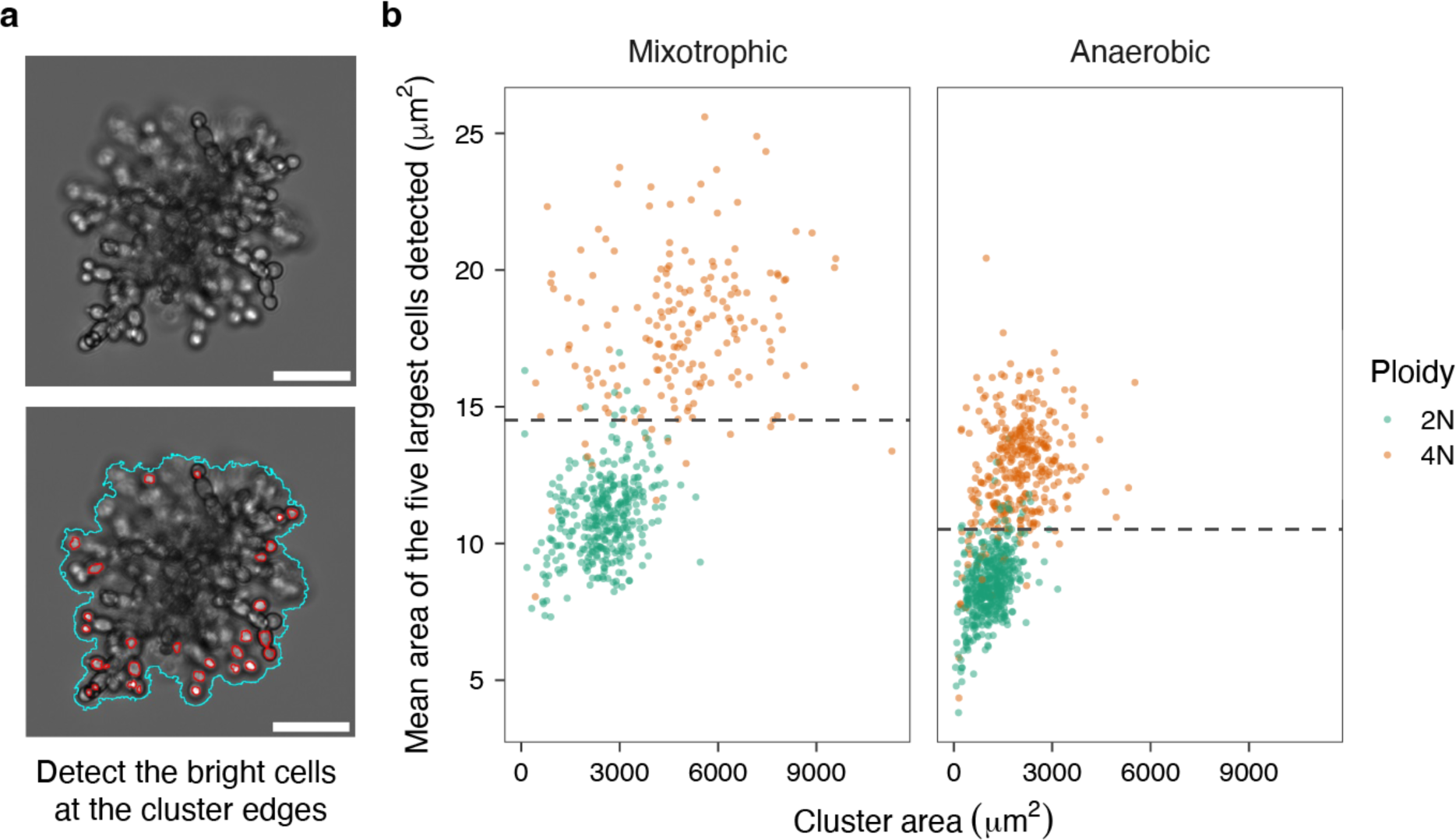
| Label-free method for distinguishing engineered diploid and tetraploid clusters in competition assays. **a**, Brightfield image of a snowflake yeast cluster (an engineered tetraploid mixotrophic cluster is shown) (top), whose bright cells in the cluster edges are detected (bottom). Scale bar, 30 µm. **b**, Mean area of the five largest cells detected in a cluster can be used to distinguish between the engineered tetraploid and diploid clusters under both mixotrophic and anaerobic conditions, with the dashed line indicating the manually-chosen decision boundary.

**Extended Data Fig. 7.**
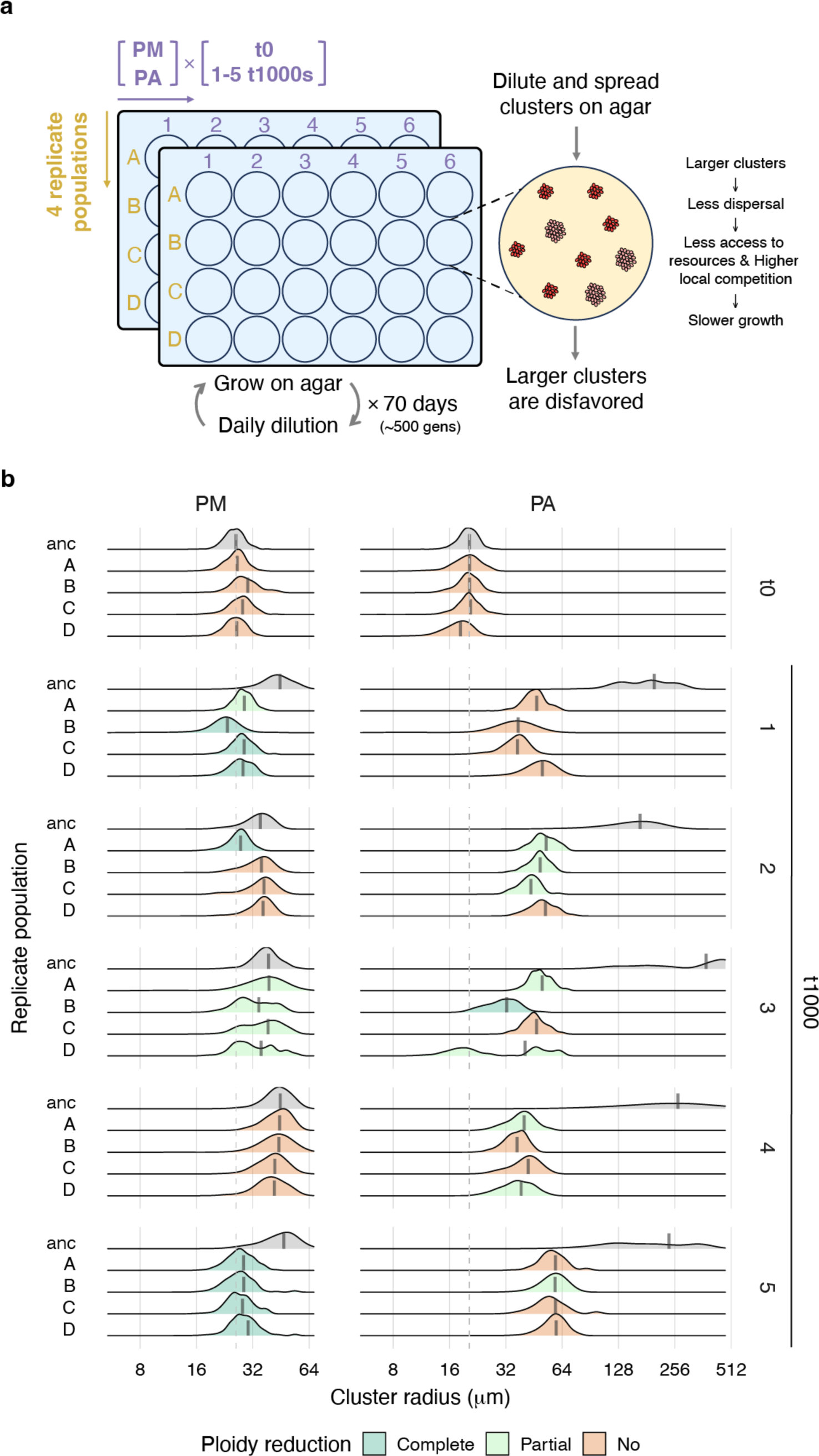
| Experimental evolution of the MuLTEE ancestors and t1000 isolates with selection against larger size. **a**, Experimental setup. We evolved PM/PA t0 and PM/PA1-5 t1000 isolates, with four replicate populations (A, B, C, D), under selection against larger size for 70 days (∼500 generations) by growing them on agar in 24-well plates with daily dilution. **b**, Distributions of cluster radius (weighted by cluster volume) in the ancestral (“anc”) and evolved populations (on average, n = 406 clusters measured per population). Vertical thick solid line, biomass-weighted mean cluster radius of each population. Vertical dashed line, biomass-weighted mean cluster radius of PM/PA t0. Color code for the level of ploidy reduction in each population is the same as that in **Fig. 3i**, and the ancestral populations are colored in gray.

**Extended Data Fig. 8.**
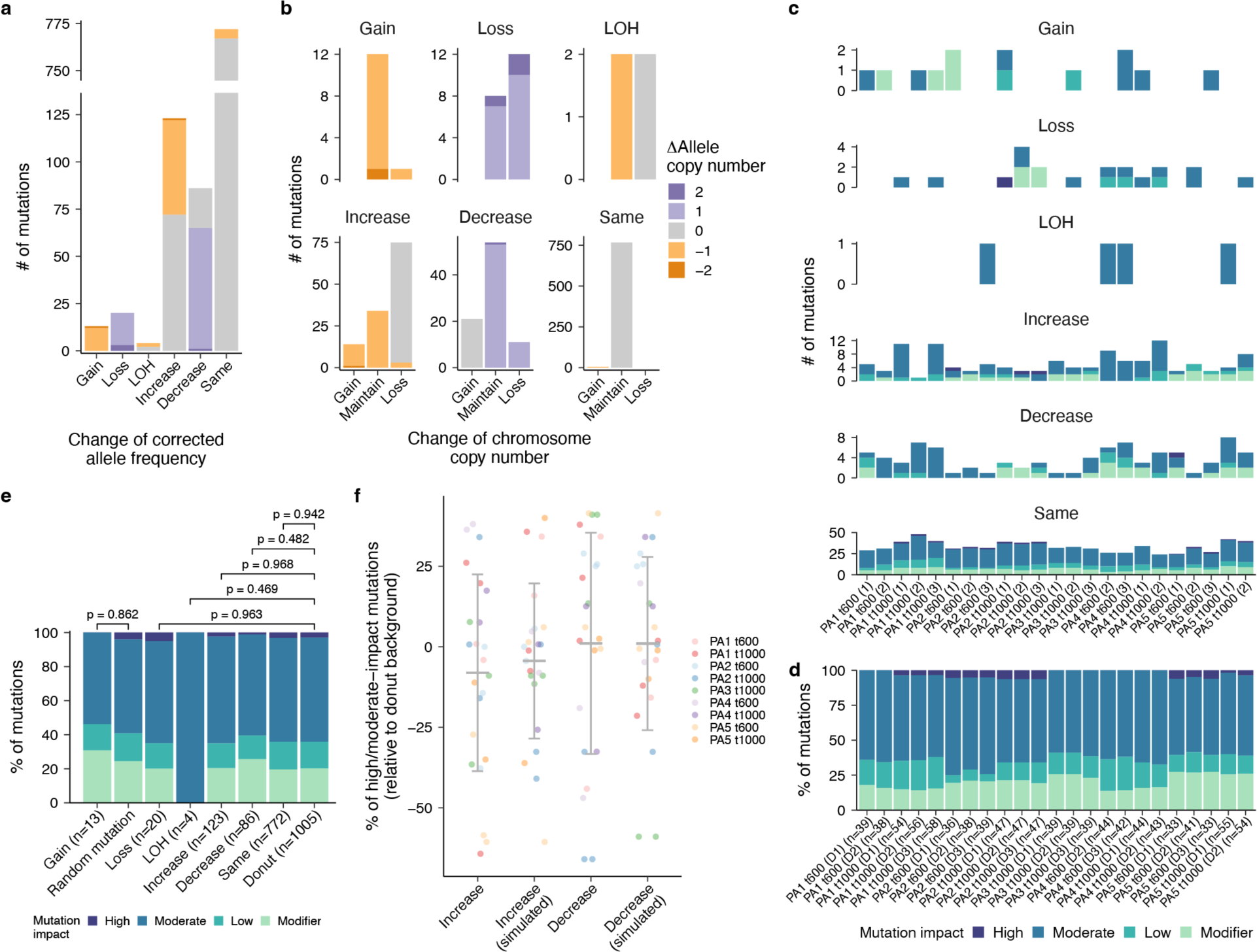
| Point mutation changes in donut-to-spread transitions. Two donut-to-spread transitions with near-triploidization were excluded, and mutation allele frequency refers to the corrected allele frequency, calculated by dividing estimated allele copy number with copy number of the chromosome that carries the mutation. LOH, loss of heterozygosity. **a,b**, Number of mutations in each category of change in allele frequency in all donut-to-spread transitions combined, colored by change in allele copy number (**a**), and how change in allele frequency is associated with change in chromosome copy number (**b**). **a,b** share the color code. **c**, Number of mutations in each category of change in allele frequency in each donut-to-spread transition, colored by mutation impact. **d**, Percentage of mutations in each mutation impact category in each donut background, whose total number of mutations is indicated in the brackets. **c,d** share the color code. **e**, Comparison of the distribution of mutation impacts, between the mutations that were gained in all donut-to-spread transitions combined and the mutations randomly introduced into yeast genome, as well as between the mutations that underwent loss, LOH, increase, decrease, or maintenance in terms of allele frequency in all donut-to-spread transitions combined and the mutations in all donut backgrounds combined. Number of mutations is indicated in the brackets. *P* values were calculated by chi-squared test. **f**, For each donut-to-spread transition, the percentage of high/moderate-impact mutations in the mutations that increased or decreased in allele frequency is on average not significantly larger than the percentage of high/moderate-impact mutations in the donut background (for increase and decrease, respectively, *P* = 0.891 and 0.442, *t*_22_ = −1.27 and 0.147, one-tailed one-sample t-test), and is largely explained by random sampling of mutations in the donut background (simulation with random seed = 1). Values are mean ± s.d. (n = 23 donut-to-spread transitions).

## REFERENCES

1 Otto, S. P. The evolutionary consequences of polyploidy. Cell 131, 452–462, doi:10.1016/j.cell.2007.10.022 (2007).

2 Selmecki, A. M. et al. Polyploidy can drive rapid adaptation in yeast. Nature 519, 349–352, doi:10.1038/nature14187 (2015).

3 Van de Peer, Y., Mizrachi, E. & Marchal, K. The evolutionary significance of polyploidy. Nat Rev Genet 18, 411–424, doi:10.1038/nrg.2017.26 (2017).

4 Vittoria, M. A., Quinton, R. J. & Ganem, N. J. Whole-genome doubling in tissues and tumors. Trends Genet 39, 954–967, doi:10.1016/j.tig.2023.08.004 (2023).

5 Comai, L. The advantages and disadvantages of being polyploid. Nat Rev Genet 6, 836–846, doi:10.1038/nrg1711 (2005).

6 Storchova, Z. et al. Genome-wide genetic analysis of polyploidy in yeast. Nature 443, 541–547, doi:10.1038/nature05178 (2006).

7 Buggs, R. J. et al. Rapid, repeated, and clustered loss of duplicate genes in allopolyploid plant populations of independent origin. Curr Biol 22, 248–252, doi:10.1016/j.cub.2011.12.027 (2012).

8 Bozdag, G. O. et al. De novo evolution of macroscopic multicellularity. Nature 617, 747–754, doi:10.1038/s41586-023-06052-1 (2023).

9 Fox, D. T., Soltis, D. E., Soltis, P. S., Ashman, T. L. & Van de Peer, Y. Polyploidy: A Biological Force From Cells to Ecosystems. Trends Cell Biol 30, 688–694, doi:10.1016/j.tcb.2020.06.006 (2020).

10 Mayrose, I., Zhan, S. H., Rothfels, C. J., Magnuson-Ford, K., Barker, M. S., Rieseberg, L. H., Otto, S. P. Recently formed polyploid plants diversify at lower rates. Science 333, 1257–1257 (2011).

11 Levin, D. A. Why polyploid exceptionalism is not accompanied by reduced extinction rates. Plant Systematics and Evolution 305, 1–11, doi:10.1007/s00606-018-1552-x (2018).

12 Clo, J. & Kolar, F. Short- and long-term consequences of genome doubling: a meta-analysis. Am J Bot 108, 2315–2322, doi:10.1002/ajb2.1759 (2021).

13 Gerstein, A. C., Chun, H. J., Grant, A. & Otto, S. P. Genomic convergence toward diploidy in Saccharomyces cerevisiae. PLoS Genet 2, e145, doi:10.1371/journal.pgen.0020145 (2006).

14 Gerstein, A. C. & Sharp, N. P. The population genetics of ploidy change in unicellular fungi. FEMS Microbiol Rev 45, doi:10.1093/femsre/fuab006 (2021).

15 Todd, R. T., Forche, A., Selmecki, A. Ploidy variation in fungi: polyploidy, aneuploidy, and genome evolution. Microbiology spectrum 5, 5–4, doi:10.1128/microbiolspec.FUNK (2017).

16 Lu, Y. J., Swamy, K. B. & Leu, J. Y. Experimental Evolution Reveals Interplay between Sch9 and Polyploid Stability in Yeast. PLoS Genet 12, e1006409, doi:10.1371/journal.pgen.1006409 (2016).

17 Bomblies, K. When everything changes at once: finding a new normal after genome duplication. Proc Biol Sci 287, 20202154, doi:10.1098/rspb.2020.2154 (2020).

18 Doyle, J. J. & Coate, J. E. Polyploidy, the Nucleotype, and Novelty: The Impact of Genome Doubling on the Biology of the Cell. International Journal of Plant Sciences 180, 1–52, doi:10.1086/700636 (2019).

19 Levin, D. A. Polyploidy and novelty in flowering plants. The American Naturalist 122, 1–25 (1983).

20 Mortier, F. et al. Understanding polyploid establishment: temporary persistence or stable coexistence? Oikos, doi:10.1111/oik.09929 (2024).

21 Van de Peer, Y., Ashman, T. L., Soltis, P. S. & Soltis, D. E. Polyploidy: an evolutionary and ecological force in stressful times. Plant Cell 33, 11–26, doi:10.1093/plcell/koaa015 (2021).

22 Storchova, Z. Ploidy changes and genome stability in yeast. Yeast 31, 421–430, doi:10.1002/yea.3037 (2014).

23 Gerstein, A. C. et al. Polyploid titan cells produce haploid and aneuploid progeny to promote stress adaptation. mBio 6, e01340–01315, doi:10.1128/mBio.01340-15 (2015).

24 Hirakawa, M. P., Chyou, D. E., Huang, D., Slan, A. R. & Bennett, R. J. Parasex Generates Phenotypic Diversity de Novo and Impacts Drug Resistance and Virulence in Candida albicans. Genetics 207, 1195–1211, doi:10.1534/genetics.117.300295 (2017).

25 Scott, A. L., Richmond, P. A., Dowell, R. D. & Selmecki, A. M. The Influence of Polyploidy on the Evolution of Yeast Grown in a Sub-Optimal Carbon Source. Mol Biol Evol 34, 2690–2703, doi:10.1093/molbev/msx205 (2017).

26 Bielski, C. M. et al. Genome doubling shapes the evolution and prognosis of advanced cancers. Nat Genet 50, 1189–1195, doi:10.1038/s41588-018-0165-1 (2018).

27 Fujiwara, T. et al. Cytokinesis failure generating tetraploids promotes tumorigenesis in p53-null cells. Nature 437, 1043–1047, doi:10.1038/nature04217 (2005).

28 Ratcliff, W. C., Fankhauser, J. D., Rogers, D. W., Greig, D. & Travisano, M. Origins of multicellular evolvability in snowflake yeast. Nat Commun 6, 6102, doi:10.1038/ncomms7102 (2015).

29 Jacobeen, S. et al. Cellular packing, mechanical stress and the evolution of multicellularity. Nat Phys 14, 286–290, doi:10.1038/s41567-017-0002-y (2018).

30 Bozdag, G. O., Libby, E., Pineau, R., Reinhard, C. T. & Ratcliff, W. C. Oxygen suppression of macroscopic multicellularity. Nat Commun 12, 2838, doi:10.1038/s41467-021-23104-0 (2021).

31 Day, T. C. et al. Morphological Entanglement in Living Systems. Physical Review X 14, doi:10.1103/PhysRevX.14.011008 (2024).

32 Day, T. C. et al. Cellular organization in lab-evolved and extant multicellular species obeys a maximum entropy law. Elife 11, doi:10.7554/eLife.72707 (2022).

33 Harrison, B. D. et al. A tetraploid intermediate precedes aneuploid formation in yeasts exposed to fluconazole. PLoS Biol 12, e1001815, doi:10.1371/journal.pbio.1001815 (2014).

34 Ramsey, J. S., D. W. Pathways, mechanisms, and rates of polyploid formation in flowering plants. Annual review of ecology and systematics 29, 467–501 (1998).

35 Gerstein, A. C., McBride, R. M. & Otto, S. P. Ploidy reduction in Saccharomyces cerevisiae. Biol Lett 4, 91–94, doi:10.1098/rsbl.2007.0476 (2008).

36 Voordeckers, K. et al. Adaptation to High Ethanol Reveals Complex Evolutionary Pathways. PLoS Genet 11, e1005635, doi:10.1371/journal.pgen.1005635 (2015).

37 Galitski, T., Saldanha, A. J., Styles, C. A., Lander, E. S., Fink, G. R. Ploidy regulation of gene expression. Science 285, 251–254 (1999).

38 Rebolleda-Gomez, M. & Travisano, M. The Cost of Being Big: Local Competition, Importance of Dispersal, and Experimental Evolution of Reversal to Unicellularity. Am Nat 192, 731–744, doi:10.1086/700095 (2018).

39 Gilchrist, C. & Stelkens, R. Aneuploidy in yeast: Segregation error or adaptation mechanism? Yeast 36, 525–539, doi:10.1002/yea.3427 (2019).

40 Vande Zande, P., Zhou, X. & Selmecki, A. The Dynamic Fungal Genome: Polyploidy, Aneuploidy and Copy Number Variation in Response to Stress. Annu Rev Microbiol 77, 341–361, doi:10.1146/annurev-micro-041320-112443 (2023).

41 O’Donnell, S. et al. Telomere-to-telomere assemblies of 142 strains characterize the genome structural landscape in Saccharomyces cerevisiae. Nat Genet 55, 1390–1399, doi:10.1038/s41588-023-01459-y (2023).

42 Anders, K. R. et al. A strategy for constructing aneuploid yeast strains by transient nondisjunction of a target chromosome. BMC Genet 10, 36, doi:10.1186/1471-2156-10-36 (2009).

43 Reid, R. J. et al. Chromosome-scale genetic mapping using a set of 16 conditionally stable Saccharomyces cerevisiae chromosomes. Genetics 180, 1799–1808, doi:10.1534/genetics.108.087999 (2008).

44 Tan, Z. et al. Aneuploidy underlies a multicellular phenotypic switch. Proc Natl Acad Sci U S A 110, 12367–12372, doi:10.1073/pnas.1301047110 (2013).

45 De Chiara, M. et al. Domestication reprogrammed the budding yeast life cycle. Nat Ecol Evol 6, 448–460, doi:10.1038/s41559-022-01671-9 (2022).

46 Albertin, W. et al. Evidence for autotetraploidy associated with reproductive isolation in Saccharomyces cerevisiae: towards a new domesticated species. J Evol Biol 22, 2157–2170, doi:10.1111/j.1420-9101.2009.01828.x (2009).

47 Yona, A. H. et al. Chromosomal duplication is a transient evolutionary solution to stress. Proc Natl Acad Sci U S A 109, 21010–21015, doi:10.1073/pnas.1211150109 (2012).

48 Schranz, M. E., Mohammadin, S. & Edger, P. P. Ancient whole genome duplications, novelty and diversification: the WGD Radiation Lag-Time Model. Curr Opin Plant Biol 15, 147–153, doi:10.1016/j.pbi.2012.03.011 (2012).

49 Li, R. & Zhu, J. Effects of aneuploidy on cell behaviour and function. Nat Rev Mol Cell Biol 23, 250–265, doi:10.1038/s41580-021-00436-9 (2022).

50 Pachitariu, M. & Stringer, C. Cellpose 2.0: how to train your own model. Nat Methods 19, 1634–1641, doi:10.1038/s41592-022-01663-4 (2022).

51 Cherry, J. M. et al. Saccharomyces Genome Database: the genomics resource of budding yeast. Nucleic Acids Res 40, D700–705, doi:10.1093/nar/gkr1029 (2012).

52. Li, H. Aligning sequence reads, clone sequences and assembly contigs with BWA-MEM. arXiv 1303, 10.48550/arXiv.1303.3997 (2013).

53 Van der Auwera, G. A. & O’Connor, B. D. Genomics in the cloud: using Docker, GATK, and WDL in Terra. (O’Reilly Media, 2020).

54 Danecek, P. et al. The variant call format and VCFtools. Bioinformatics 27, 2156–2158, doi:10.1093/bioinformatics/btr330 (2011).

55 Danecek, P. et al. Twelve years of SAMtools and BCFtools. Gigascience 10, doi:10.1093/gigascience/giab008 (2021).

56 Cingolani, P. et al. A program for annotating and predicting the effects of single nucleotide polymorphisms, SnpEff: SNPs in the genome of Drosophila melanogaster strain w1118; iso-2; iso-3. Fly (Austin) 6, 80–92, doi:10.4161/fly.19695 (2012).

57 Todd, R. T., Braverman, A. L. & Selmecki, A. Flow Cytometry Analysis of Fungal Ploidy. Curr Protoc Microbiol 50, e58, doi:10.1002/cpmc.58 (2018).

