## Supplementary File 1 for "Whole-genome duplication in the Multicellularity Long Term Evolution Experiment"

### **Analysis of point mutation changes in donut-to-spread transitions**

Besides karyotype changes, we also examined the potential roles of changes in point mutations (single- or multi-nucleotide substitutions or indels) in donut-to-spread transitions, as our previous work^1^ implicated the role of evolved point mutations in the origin of macroscopic size. For each point mutation, we first estimated its allele copy number based on its allele frequency and chromosome copy number, and then corrected its allele frequency estimate using its allele copy number and chromosome copy number (Methods). Based on the corrected allele frequencies, we classified point mutation changes into six categories: gain, loss, loss of heterozygosity (LOH, i.e., reach 100% allele frequency from below 100%, essentially loss of ancestral allele), increase and decrease in allele frequency, and retain the same allele frequency. We found that when the allele copy number of a point mutation changed in the donut-to-spread transitions, it mostly changed by one copy (Extended Data Fig. 8a). This implies that most mutation gains were likely spontaneous mutations that occur in one of multiple chromosome copies, and most mutation losses occurred to mutations carried by only one of multiple chromosome copies (Extended Data Fig. 8a). Mutation allele frequency can, however, change without changing its allele copy number (Extended Data Fig. 8a), and these cases were all associated with chromosome gains or losses (likely in the chromosome copies that do not carry the mutation) (Extended Data Fig. 8b). When a mutation allele frequency change involves changing its allele copy number, it can occur with chromosome gain or loss (likely in the chromosome copies that carry the mutation) (Extended Data Fig. 8b), or without chromosome copy number change (Extended Data Fig. 8b), likely involving gene conversion between the mutant allele and the ancestral allele on different chromosome copies. The latter scenario (without chromosome copy number change) occurred frequently, making up 98 of 233 (~42%) allele frequency changes in pre-existing mutations (i.e., excluding mutation gains).

To explore the functional importance of point mutation changes in donut-to-spread transitions, we followed the classification system in SnpEff^2^ and classified the potential impacts of observed point mutations into high (nonsense and frameshift variants), moderate (missense and in-frame deletion variants), low (synonymous and splice region variants), and modifier (upstream gene variants). We found that each donut-to-spread transition had allele frequency changes in an average of 11 point mutations, most of which are increases or decreases in allele frequencies and have high or moderate impacts (Extended Data Fig. 8c). If point mutation changes are critically important in the donut-to-spread transitions, we would expect high/moderate-impact mutations to be enriched in the mutations that changed allele frequencies, in a similar vein to how high dN/dS ratio signifies positive selection in evolution. However, the impact distribution of all changed mutations in donut-to-spread transitions were not statistically significantly different from what is expected from random new mutations or random sampling of pre-existing mutations in all donut backgrounds (Extended Data Fig. 8d,e). For each donut-to-spread transition, the percentage of high- and moderate-impact mutations in mutations with allele frequency increases or decreases could also be largely explained by random sampling of mutations in the corresponding donut background (Extended Data Fig. 8d,f). Nevertheless, these do not rule out the possibility that changes in some point mutations could have contributed to the loss of macroscopic size. Indeed, the first donut-spread pair from the PA1 t600 isolate was not associated with changes in karyotype (Fig. 4h) but in point mutations (Extended Data Fig. 8c), suggesting the potential role of point mutations in this case. However, overall, we found no strong statistical evidence that point mutations played a systematic role in the rapid loss of macroscopic size during the donut-to-spread transitions, while we found very strong evidence for the critical role of aneuploidy in this process (Fig. 4g-o).

**References**

1	Bozdag, G. O. *et al.* De novo evolution of macroscopic multicellularity. *Nature* **617**, 747-754, doi:10.1038/s41586-023-06052-1 (2023).

2	Cingolani, P. *et al.* A program for annotating and predicting the effects of single nucleotide polymorphisms, SnpEff: SNPs in the genome of Drosophila melanogaster strain w1118; iso-2; iso-3. *Fly (Austin)* **6**, 80-92, doi:10.4161/fly.19695 (2012).
